# Neurochemical Features and Glycan Patterns in the Wapiti Vomeronasal Pathway

**DOI:** 10.1101/2025.06.25.661586

**Authors:** Antía Martínez Antonio, Gadea García Hernando, Esther Valderrábano Cano, José Luis Rois, Óscar Sampedro Moreira, Pablo Sanchez Quinteiro, Irene Ortiz Leal

**Affiliations:** Department of Anatomy, Animal Production and Clinical Veterinary Sciences. Faculty of Veterinary. University of Santiago de Compostela. Lugo 27002, Spain; Marcelle Nature Park, Outeiro de Rei, Lugo, Spain

**Keywords:** Vomeronasal system, Cervus canadensis, immunohistochemistry, lectin histochemistry, accessory olfactory bulb, chemical communication, wild ruminants

## Abstract

The vomeronasal system (VNS) plays a central role in mammalian chemical communication, mediating critical social and reproductive behaviors. In the wapiti (*Cervus canadensis*), a cervid species with complex social structures and pronounced chemical signaling during the rut, the VNS had not been previously characterized. This study provides the first comprehensive anatomical and neurochemical analysis of the VNS in wapiti using histological, lectin-histochemical, and immunohistochemical techniques. The vomeronasal organ (VNO) exhibited clear rostrocaudal differentiation, with distinct sensory and respiratory epithelia, a complex glandular distribution, and region-specific expression of neural markers. Lectin binding patterns confirmed functional compartmentalization along the epithelium, and immunoreactivity for markers such as OMP, PGP9.5, CR, and G-protein subunits (Gαi2, Gγ8, and Gα0) revealed detailed molecular organization. Notably, Gα0-positive neurons in the epithelium did not project to the accessory olfactory bulb (AOB), suggesting alternative targets, possibly within transitional zones. The AOB showed all canonical layers, including well-defined glomeruli and expression of markers such as calbindin, CR, GFAP, and LEA lectin. Novel findings include the presence of large white matter tracts and region-specific lectin distribution. Confocal double immunofluorescence and autofluorescence imaging were also employed, allowing high-resolution visualization of neuroepithelial architecture and glomerular domains. Altogether, our results demonstrate that the vomeronasal system in wapiti is highly developed and functionally specialized. These findings contribute to a better understanding of chemosensory communication in wild ungulates and provide a comparative framework for future studies in cervid behavior, reproduction, and conservation.

## Introduction

The wapiti (*Cervus canadensis*), commonly known as North American elk, is one of the largest living cervid species, surpassed in body size only by the moose (*Alces alces*) (Geist, 1998). Native to vast temperate and boreal regions of North America and parts of eastern Asia, wapiti populations exhibit pronounced social structures, seasonal reproductive cycles, and highly ritualized behaviors. Males, particularly during the rut, demonstrate aggressive and reproductive displays that rely heavily on chemical cues: they urinate on their own bodies, wallow in scent-marked soil, and engage in vocalizations and physical confrontations to establish dominance and attract females (Toweill and Thomas, 2002). These behavioral strategies strongly suggest the existence of a sophisticated system of chemosensory communication that integrates social, sexual, and environmental information. Among vertebrates, chemosensory communication is essential for detecting conspecifics, marking territory, locating food, and identifying reproductive partners (Wilson, 1970; Allen et al., 2014). This form of communication is mediated by the olfactory system, which includes the main olfactory system and the vomeronasal system (VNS) (Barrios et al., 2014). While the main olfactory system is traditionally associated with the detection of a broad range of volatile odorants, the VNS—also referred to as the accessory olfactory system—is highly specialized for detecting semiochemicals such as pheromones and kairomones (Brennan and Zufall, 2006; Ruiz-Rubio et al., 2024). These substances are crucial in modulating social and reproductive behaviors across many mammalian species and can trigger both immediate behavioral responses and long-term physiological changes (Wyatt, 2013; Brennan, 2018).

In ruminants, pheromones have been shown to influence a wide array of physiological and behavioral processes, including sexual attraction, mother– offspring bonding, estrus signaling, induction and synchronization, puberty onset, postpartum anestrus, hormonal activation, and penile erection (Archunan et al., 2014; Rajanarayanan and Archunan, 2004). Moreover, semiochemicals play a role in stress regulation mechanisms in mammals, and it has been proposed that they may help mitigate male aggression in the context of reintroduction programs by modulating social behavior (Archer et al., 2022). Although these mechanisms remain unexplored in wapiti, studies in other ruminants, such as the dama gazelle, have highlighted the potential role of semiochemical communication in regulating both reproductive behavior and stress-related social dynamics (Torres et al., 2023a).

Anatomically, the vomeronasal system consists of the vomeronasal organ (VNO), the vomeronasal nerve, and the accessory olfactory bulb (AOB) (Torres et al., 2022). Vomeronasal sensory neurons express specific families of receptors—V1Rs (Boschat et al., 2002), V2Rs (Ortiz-Leal et al., 2024), and formyl peptide receptors (FPRs) (Ackels et al., 2014)—and project their axons to the AOB, where the first synaptic integration of the chemical signal occurs (Dulac and Axel, 1995; Tirindelli, 2021). This information is then conveyed to deeper regions of the limbic system, such as the hypothalamus and amygdala, which are involved in emotional and endocrine regulation (Larriva-Sahd et al., 1993; Gutiérrez-Castellanos et al., 2014). The close association between the VNS and the hypothalamic–pituitary axis highlights its relevance in coordinating behavioral, hormonal, and physiological responses to external chemical stimuli (Salazar et al., 2016; Torres et al., 2023b).

Wapiti are phylogenetically classified within the order Artiodactyla, suborder Ruminantia, and infraorder Pecora. Within this group, they belong to the family Cervidae, which also includes red deer, moose, and roe deer. The infraorder additionally comprises Bovidae, Antilocapridae (pronghorns), Giraffidae, and Moschidae (musk deer). Comparative research on the vomeronasal system across these groups has revealed notable anatomical and functional diversity. In giraffes and musk deer, for example, the VNO has been histologically characterized, with recent studies documenting both its structure and glycoconjugate composition (Kondoh et al., 2017b; Hart and Hart, 2023; Kondoh et al., 2020). Although more extensive studies have been conducted in bovids, most of them have focused on domestic species such as cattle (*Bos taurus coreanae*) (Jacobs et al., 1981; Taniguchi and Mikami, 1985; Adams, 1986; Salazar et al., 2008; Jang et al., 2021), goats (*Capra aegagrus hircus*) (Ladewig and Hart, 1980; Park et al., 2013, 2013), and sheep (*Ovis orientalis aries*) (Kratzing, 1971; Salazar et al., 1998, 2000, 2003, 2007; Ibrahim et al., 2013; Barrios et al., 2014; Ibrahim, 2018), while relatively little attention has been paid to wild bovids.

Within Cervidae, the vomeronasal system has been shown to be well developed and functional in several species, including reindeer (*Rangifer tarandus*) (Bertmar, 1981), moose (*Alces alces*) (Vedin et al., 2010), sika deer (*Cervus nippon*) (Matsubara et al., 2019), and Korean roe deer (*Capreolus pygargus*) (Park et al., 2014; Shin et al., 2017). Altogether, these studies highlight the need for further anatomical and neurochemical investigation of the vomeronasal system in wild cervids, particularly in ecologically and behaviorally complex species such as wapiti.

Despite the taxonomic and ecological importance of the wapiti, to our knowledge no detailed morphological or neurochemical studies of its vomeronasal system have been published to date. This gap is particularly notable given the pronounced behavioral reliance of wapiti on chemical cues during social and reproductive interactions. In this context, the present study was designed to provide a comprehensive morphofunctional characterization of the vomeronasal system in *Cervus canadensis*. Using an integrative approach combining histological, lectin-histochemical, and immunohistochemical techniques, we investigated the anatomical organization and neurochemical features of the VNO and the olfactory bulbs—both main and accessory. Lectin histochemistry was employed to visualize patterns of glycosylation, which are indicative of functional compartmentalization within the sensory epithelium and neural structures (Ortiz-Leal et al., 2022b). Immunohistochemical markers were used to identify specific neuronal populations and their projection patterns, allowing us to assess the degree of structural and functional differentiation along the rostrocaudal axis of the VNO and the AOB (Torres et al., 2021).

By documenting the histological features and molecular signatures of the vomeronasal system in a non-domestic cervid species, our findings aim to expand the current understanding of interspecific variability in chemosensory systems among ruminants. Moreover, this study establishes a foundational anatomical and neurochemical reference for future research on the role of semiochemicals in wapiti behavior and reproduction. Such knowledge could have practical implications for wildlife conservation programs, assisted reproduction, and the management of wild cervid populations in both natural and captive settings.

## Methods

### Origin of the samples

This study was conducted using a single specimen of wapiti (W) from the Marcelle Nature Park, located in the municipality of Outeiro de Rei, Lugo province (Galicia, Spain). The animal was a 20-year-old female that died of natural causes associated with physiological aging, without any apparent signs of disease. The entire body was not available; only the head was received at the dissection laboratory of the Department of Anatomy. From this specimen, the vomeronasal organs embedded in the nasal septum and the complete brain were also obtained. For the purposes of the present study, only the anterior portion of the brain, including the main and accessory olfactory bulbs, was analyzed. Finally, the rostral region of the palate, which includes the incisive papilla, was dissected and removed.

All specimens obtained from the head dissection were fixed in Bouin’s solution for 24 hours and subsequently stored in 70% ethanol until paraffin embedding. Bouin’s fixative was selected due to its excellent capacity to preserve tissue morphology and architecture. Additionally, it provides strong contrast for histological staining and generally eliminates the need for antigen retrieval in immunohistochemistry. The fixative consists of picric acid, formaldehyde, and acetic acid.

### Sample extraction

#### Olfactory bulbs (OB)

To extract the brain, the skin and underlying musculature of the skull were first removed. Then, incisions were made in the bones forming the dorsal, lateral, and caudal walls of the cranial cavity (frontal, temporal, parietal, and occipital bones), taking particular care around the ethmoidal fossa, the region housing the olfactory bulbs. Using a rongeur forceps, the cranial vault was lifted, thereby exposing the olfactory bulbs and cerebral hemispheres. The entire brain was immersed in Bouin’s fixative for 24 hours, during which it acquired sufficient rigidity to allow for complete dissection. The olfactory bulbs were then carefully separated from the multiple olfactory nerve bundles anchoring them to the cribriform plate of the ethmoid bone.

#### Vomeronasal organ (VNO)

Accessing the VNO required exposure of the vomer bone. Therefore, the nasal bones and lateral portions of the maxillary bones were first removed. The vomer region was then delimited, and the bone was extracted along with the vomeronasal organs. Immediate fixation in Bouin’s solution followed. After fixation, the vomeronasal organs were dissected out over the following days using blunt dissection techniques with fine forceps. To facilitate subsequent histological sectioning, a decalcification process was required for those specimens containing bone tissue (incisive papilla and palatal region). Decalcification was performed using Osteomoll solution (Sigma) under continuous agitation for 14 days.

### Paraffin Embedding

Paraffin embedding facilitates sectioning of the samples using a rotary microtome. This process requires prior dehydration of the tissues, which is achieved by immersing the samples in a graded ethanol series. The steps followed were as follows: 70% ethanol (2 hours), 90% ethanol (1 hour), 96% ethanol (2 hours), and three changes of 100% ethanol (1 hour each). Subsequently, tissue clearing was initiated with a mixture of xylene and ethanol (1 hour and 15 minutes), followed by two immersions in pure xylene (1 hour and 30 minutes, then 30 minutes). The process concluded with two consecutive paraffin baths of 2 hours each. Finally, the sample was placed in a mold and filled with hot liquid paraffin to form a solid block upon cooling. To accelerate the solidification process, the mold was placed on a cold plate.

### Microtome Sectioning

For sectioning the paraffin blocks, a Leica Reichert Jung Autocut 2055 microtome was used. This device allows sectioning at variable thicknesses depending on the tissue type. For nervous tissue, sections were cut at 8 µm thickness due to its low cellular density, which allows better visualization of cellular processes. Due to the longitudinal size of the vomeronasal organ (15 cm), it was necessary to divide the organ into different regions by means of transverse sections in order to facilitate paraffin embedding and microtome sectioning. All segments were included in the study. The organ was divided into 15 segments, each approximately 1 cm in length, thereby yielding more manageable portions. From each of the 15 regions, histological sections were obtained for various stains, and ribbon samples were collected and preserved for subsequent immunohistochemical and lectin histochemistry analyses.

### General histological stains

For the realization of this study, we have used the following stains:

#### Hematoxylin-eosin (H-E)

This stain was selected for the general observation of the various structures, as it provides clear visualization of cellular and tissue morphology. Hematoxylin is a basic dye that stains cell nuclei, while eosin is an acidic dye that stains the cytoplasm.

#### Nissl Staining

This technique is used exclusively for nervous tissue, as it allows clear visualization of glial cells and neuronal somata along with the initial segments of their processes. It enables the assessment of cell location, distribution, and density within nervous tissue. The staining relies on cresyl violet, which binds to nucleic acid-rich components such as free cytoplasmic ribosomes, rough endoplasmic reticulum, and the nucleus.

#### PAS

Periodic Acid–Schiff (PAS) staining is a special histological technique used to detect polysaccharides such as glycogen, mucins, and glycoproteins. The principle of the stain is based on the oxidation of carbohydrate groups in the tissue by periodic acid, producing aldehyde groups. These newly formed aldehydes then react with Schiff reagent, resulting in a characteristic magenta coloration. The protocol followed is detailed in depth in Villamayor et al. (2021).

#### Alcian blue (AB)

This basic dye is used to stain acidic substances, primarily acidic mucopolysaccharides. It binds to carbohydrates and is used in combination with PAS staining to differentiate between acidic and neutral mucins. Alcian Blue is a cationic dye that binds to anionic groups of acidic mucins, producing a turquoise-blue color. This staining is particularly effective in glandular tissues, as well as in structural components such as cartilage and bone. The detailed protocol applied is specified in Ruiz-Rubio et al. (2023).

#### Gallego’s Trichrome Staining

This is a compound staining technique that differentially stains various tissue components. Ferric hematoxylin: stains cell nuclei dark blue or black. Acid fuchsin: stains muscle fibers and cytoplasm bright red. Light green or aniline blue: stains collagen green. The protocol is detailed in depth in Ortiz-Leal et al. (Ortiz-Leal et al., 2024).

### Immunohistochemical staining

To investigate the expression patterns of specific neuronal markers and G protein subunits, we performed immunohistochemical analyses using an indirect peroxidase-based method. This technique relies on the high specificity of antigen–antibody binding and enables microscopic visualization of target molecules in paraffin-embedded tissue sections. Tissue sections were deparaffinized in xylene and rehydrated through a graded ethanol series into 0.1 M phosphate buffer (PB). This step aimed to reestablish physiological osmotic conditions prior to antibody incubation. To prevent nonspecific background staining caused by endogenous peroxidase activity, sections were incubated in a 3% hydrogen peroxide (H₂O₂) solution prepared in distilled water. Subsequently, nonspecific binding sites were blocked using 2% bovine serum albumin (BSA) diluted in 0.1 M phosphate buffer. A two-step immunohistochemical procedure was applied:

#### Primary antibody

Tissue sections were incubated overnight at 4 °C with the appropriate primary antibody.

#### Secondary antibody

Sections were then incubated for 30 minutes at room temperature with a peroxidase-conjugated secondary antibody polymer (CRF Anti-Polyvalent HRP Polymer; ScyTek, Cache Valley, USA).

Following primary and secondary antibody incubations, unbound antibodies were removed by successive washes in phosphate buffer (PB), with a final rinse in 0.1 M TRIS buffer. Visualization of the immunoreactive sites was achieved using 3,3′-diaminobenzidine (DAB) as the chromogen in the presence of hydrogen peroxide. The peroxidase catalyzed the oxidation of DAB, leading to a localized brown precipitate at sites of antibody binding. Sections were then rinsed in PB and distilled water and mounted with a mounting medium for light microscopy.

Positive controls included tissue known to express the target antigen, while negative controls were processed in parallel by omitting the primary antibody to confirm the specificity of the immunostaining.

### Primary antibodies

The following primary antibodies were used:

<u>Anti-Gα₀</u>: recognizes the α₀ subunit of G proteins; associated with the V2R vomeronasal transduction pathway in the VNO and AOB (Ortiz-Leal et al., 2023).

<u>Anti-Gαi₂</u>: targets the αi₂ subunit of G proteins; indicative of V1R-expressing vomeronasal pathways (Ortiz-Leal et al., 2022a).

Anti-Gγ₈: recognizes the γ₈ subunit; expressed in the olfactory and vomeronasal epithelia, associated with neurogenesis (Ryba and Tirindelli, 1995).

<u>Anti-GAP43</u>: detects growth-associated protein 43, involved in axonal growth and synaptic plasticity (Ruiz-Rubio et al., 2024).

<u>Anti-calbindin</u>: a calcium-binding protein implicated in intracellular calcium buffering in excitable cells (Porteros et al., 1995).

<u>Anti-PGP9.5</u>: neuronal marker for ubiquitin carboxyl-terminal hydrolase, widely expressed in neurons (Lundberg et al., 1988).

<u>Anti-calretinin</u>: another calcium-binding protein, found in subsets of neurons with specific physiological properties (Briñón et al., 2001).

<u>Anti-neuron-specific enolase</u>: a glycolytic enzyme used as a general neuronal marker (Takahashi et al., 1984).

Anti-MAP2: detects microtubule-associated protein 2, localized in dendrites and involved in cytoskeletal stabilization (Johnson and Jope, 1992).

### Lectin histochemical labelling

Lectin-based histochemistry is conceptually analogous to immunohistochemistry, but it does not rely on immune-derived components. Lectins are glycan-binding proteins, or agglutinins, capable of binding specific carbohydrate residues in a reversible and non-covalent manner without chemically modifying the glycoconjugates. These molecules are derived from various organisms, most commonly plants, and are named accordingly (Plendl and Sinowatz, 1998). In the present study, we employed ten lectins to characterize the glycoconjugate composition of different olfactory and vomeronasal structures. All lectins were applied to all tissue samples at dilutions optimized in preliminary trials. Their sources, carbohydrate affinities, and known specificities are as follows:

STL (*Solanum tuberosum* lectin): binds N-acetylglucosamine (GlcNAc) and its oligomers (McCurrach and Kilpatrick, 1986).

<u>LEA (*Lycopersicon esculentum* agglutinin):</u> recognizes GlcNAc-rich N-glycans (Tomiyasu et al., 2018).

<u>DBA (*Dolichos biflorus* agglutinin):</u> shows high affinity for α-GalNAc, particularly terminal residues of blood group A antigens (Chun et al., 2024).

<u>SBA (*Glycine max* agglutinin):</u> binds primarily to β-galactose (Taniguchi et al., 1993).

<u>VVA (*Vicia villosa* agglutinin):</u> targets D-galactose and N-acetylgalactosamine (GalNAc) (Shapiro et al., 1995).

<u>UEA (*Ulex europaeus* agglutinin):</u> binds terminal L-fucose; selectively labels olfactory and vomeronasal components (Kondoh et al., 2017a).

<u>ECL (*Erythrina cristagalli* lectin):</u> binds D-galactose and GlcNAc residues (Keller et al., 2022).

<u>WGA (*Triticum vulgaris* lectin):</u> GlcNAc and sialic acid (Lee et al., 2012).

<u>PHL (*Phaseolus lunatus* lectin)</u>: recognizes mannose residues, and to a lesser extent, glucose (Shin et al., 2017).

<u>LCA (*Lens culinaris* agglutinin)</u>: binds D-galactose residues (Salazar et al., 2008).

### Labeling procedure

Tissue sections were deparaffinized, rehydrated through graded ethanol series, and transferred to phosphate buffer (0.1 M, pH 7.4). To prevent nonspecific binding, slides were incubated for 30 minutes in 2% bovine serum albumin (BSA) in phosphate buffer. Lectin labeling was performed using protocols specific to each lectin’s conjugation. Biotinylated lectins were applied and incubated overnight at 4°C. On the following day, the sections were incubated for 90 minutes with an avidin–biotin–peroxidase complex (ABC Elite Kit, Vectastain, Vector Labs), which binds to the biotin moiety of the lectin. Lectin binding was visualized using a chromogenic reaction based on 3,3′-diaminobenzidine (DAB). Slides were incubated in a freshly prepared solution containing 0.05% DAB and 0.003% H₂O₂ in 0.2 M Tris-HCl buffer (pH 7.6). The peroxidase activity catalyzed the oxidation of DAB, producing an insoluble brown precipitate at the binding sites.

### Immunohistochemistry for Optical Microscopy in Free-floating sections

Tissues destined for cryosectioning underwent a cryoprotection protocol to prevent the formation of ice crystals during the freezing process. This was achieved by immersing the samples in a graded series of sucrose solutions (10%, 20%, and 30% w/v). Sections were obtained using a sliding microtome equipped with a cooling system. Tissue was cut at a thickness of 50 µm. Sections were collected in wells containing phosphate buffer (PB) and stored at 4 °C until further processing. Sections underwent immunohistochemical staining using a floating section protocol (sections were incubated in wells rather than mounted on glass slides). The steps followed were identical to those described for paraffin-embedded samples.

### Double Immunohistochemistry for Confocal Fluorescence Microscopy in Free-floating sections

Confocal microscopy was performed to visualize dual immunolabeling using monoclonal and polyclonal antibodies. The protocol followed the same steps as for floating sections, with the addition of a Triton X-100 permeabilization step (15% solution, 15 min) prior to blocking with BSA and primary antibody incubation. Secondary antibodies were conjugated to Alexa fluorochromes (NeoBiotech): Alexa Fluor 488 anti-mouse IgG and Alexa Fluor 568 anti-rabbit IgG, both diluted 1:250 in PB. Sections were incubated for 1 hour at room temperature, under agitation and protected from light. Afterward, they were washed three times in PBS (in darkness to preserve fluorescence). Nuclear counterstaining was performed using TO-PRO-3 iodide (Molecular Probes) at a dilution of 1:400.

### Autofluorescence Imaging

Autofluorescence microscopy was used to visualize endogenous fluorescence of specific structures without applying exogenous markers. This technique was applied to three cryosections of the vomeronasal organ (VNO). Transverse sections (40 µm) were obtained and collected in 0.1 M PBS (pH 7.2), then stored at 4 °C until processed as free-floating sections. A Leica TCS SPE confocal system was used, configured for dual-channel fluorescence detection (green autofluorescence and far-red fluorescence with TO-PRO-3). Laser settings included a 100 mW argon laser (488 nm) and a helium-neon laser (633 nm). Nuclear DNA staining was achieved by incubating sections with TO-PRO-3 iodide (1 µM; Molecular Probes) for 15 minutes. After three PBS washes, sections were mounted in SlowFade antifade mounting medium (Molecular Probes).

### Image Acquisition

Digital images were captured using an Olympus SC180 digital camera mounted on an Olympus BX50 microscope (Tokyo, Japan). To ensure high-definition imaging of large tissue areas, final images were created as mosaics composed of up to 100 individual microphotographs merged using the automatic stitching software PTGui (Rotterdam, Netherlands). Adobe Photoshop CS4 (Adobe Systems, San Jose, CA) was used to adjust brightness, contrast, and white balance, and to crop and standardize image dimensions. No enhancements, additions, or alterations to the image characteristics were performed.

## Results

### Macroscopic study of the VNO

Macroscopic examination of the head of the wapiti (*Cervus canadensis*) enabled clear identification of the anatomical structures involved in the chemosensory pathway associated with the vomeronasal organ (VNO). The dorsolateral view of the head provided overall orientation and allowed for a general morphological assessment of the specimen (Fig. 1A). On the ventral surface of the palate, a well-defined incisive papilla was observed (Fig. 2B), located in the anterior region of the palatal mucosa. This structure constitutes the entry point for semiochemical signals into the vomeronasal system by connecting the oral cavity with the VNO via the incisive duct. A magnified image of this region (Fig. 1C) showed the precise topographical arrangement of the incisive papilla (IP) in relation to the palatine ridges (PR) and the upper lip (UL), highlighting its accessible location and potential functional relevance during oral or facial exploratory behaviors. Following fixation and dissection of the palatal block, the bilateral incisive ducts and their continuity with the vomeronasal system were clearly identified (Fig. 1D). Finally, the medial view of the left VNO revealed its tubular morphology, along with the course of the vomeronasal nerves emerging dorsally from the organ toward the accessory olfactory bulb (Fig. 1E).

**Figure 1.**
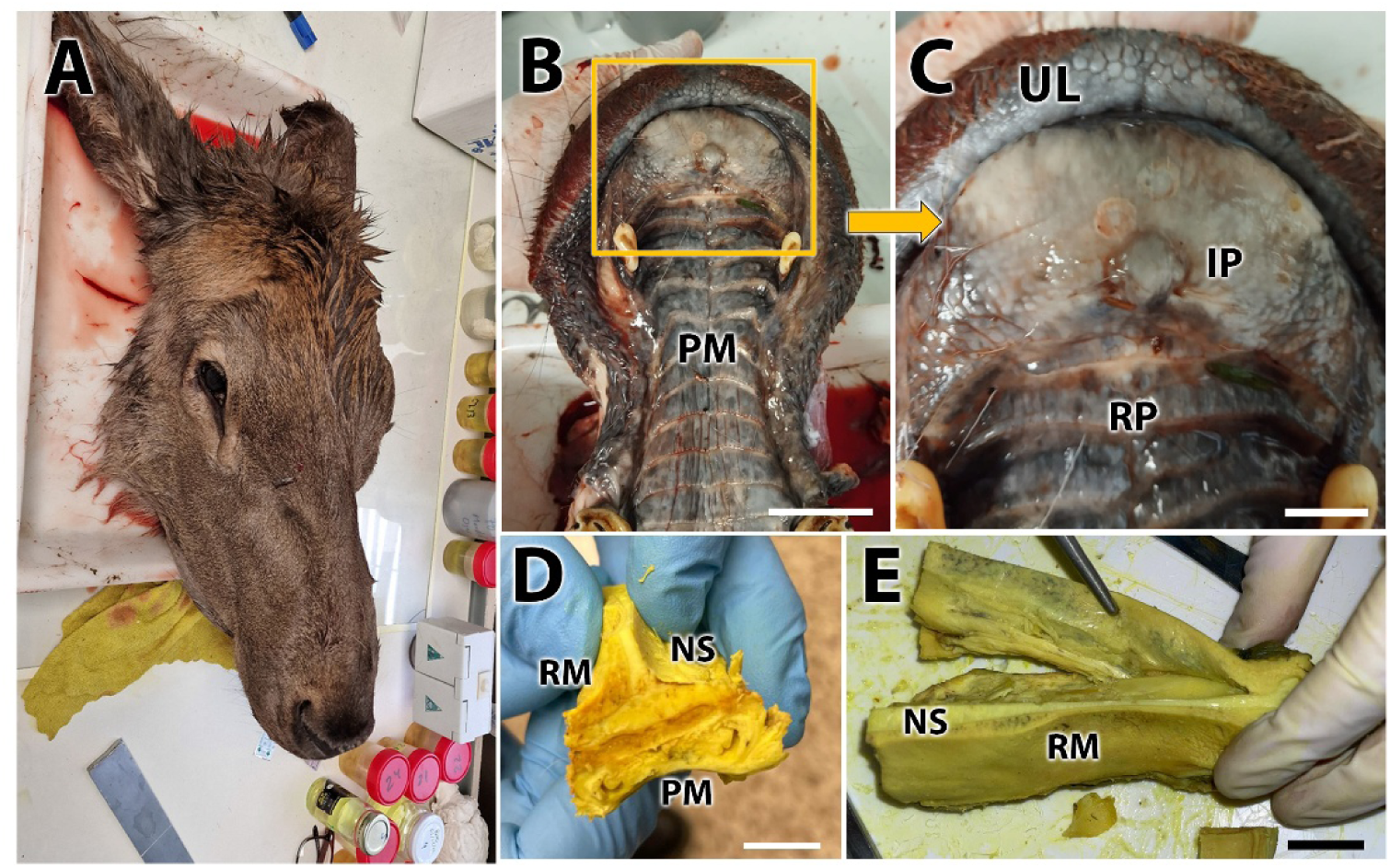
Macroscopic study of the vomeronasal system in the wapiti. **A.** Dorsolateral view of the head of the studied wapiti specimen (*Cervus canadensis*). **B.** Ventral view of the palatal mucosa showing the incisive papilla, a structure that enables the access of semiochemicals from the oral cavity to the VNO through the incisive duct. **C.** Magnified view of the area surrounding the incisive papilla (IP), illustrating its topographical relationship with the palatine ridges (PR) and the upper lip (UL). **D.** Palatal tissue block extracted after fixation and dissection, allowing identification of the incisive ducts within the palatal mucosa. **E.** Medial view of the left VNO (forceps) showing the trajectory of the vomeronasal nerves (arrowhead). Scale bars. A-E = 1 cm.

**Figure 2.**
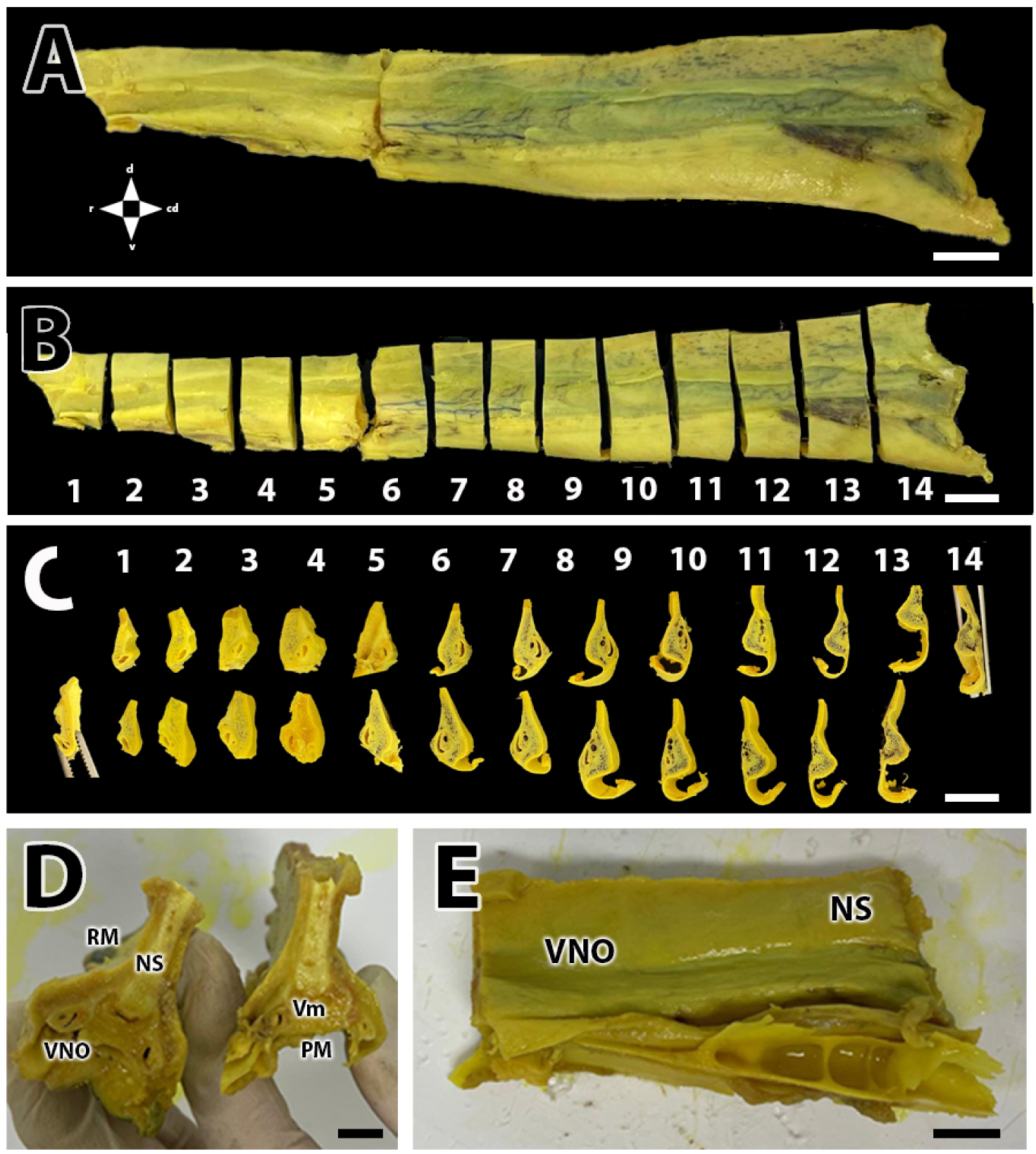
Macroscopic images of the VNO of the wapiti following its extraction and fixation in Bouin’s solution. A. General view of the entire VNO as extracted. B. Series of transverse segments of the VNO after division into 14 levels for differential analysis. Each level was approximately 1 cm in length. C. Composite view showing the rostral (top row) and caudal (bottom row) aspects of each transverse section. D. Rostral (left) and caudal (right) views of the nasal septum in transverse section, illustrating the topographical relationships of the VNO with the vomer bone (Vm), nasal septum (NS), and respiratory mucosa (RM). E. Lateral view of the caudal portion of the VNO depicted in panel A. Scale bar: (A–E) 1 cm.

After extraction and fixation of the Buoin’s liquid fixed VNO, macroscopic images were obtained to characterize its morphology and topography. The general view of the intact VNO (Fig. 2A) displayed its preserved longitudinal arrangement. The organ was then sectioned transversely into 14 consecutive levels, each approximately 1 cm in length, enabling a segmented analysis along its rostrocaudal axis (Fig. 2B). The comparative montage of transverse sections clearly revealed differences along the rostral and caudal ends of the organ (Fig. 2C). Additionally, transverse sections of the nasal septum were obtained (Fig. 2D), revealing the topographical relationships of the VNO with adjacent structures, particularly the vomer bone, the nasal septum, and the respiratory mucosa. These observations allowed anatomical contextualization of the VNO’s location within the nasal complex. A lateral view of the caudal portion of the VNO (Fig. 2E) provided a detailed depiction of the external morphology in this segment.

### Microscopic study of the VNO

To illustrate the histological organization of the VNO, a representative image from a central level of the organ was selected and stained with Gallego’s trichrome (Fig. 3). This staining technique allowed clear identification of the main components of the system, including the vomeronasal cartilage, the vomeronasal duct, and the vascular pump. Additionally, the topographical relationship between the VNO and the respiratory mucosa of the nasal cavity was evident, with the presence of associated glands and numerous blood vessels, highlighting the structural and functional complexity of this region.

**Figure 3.**
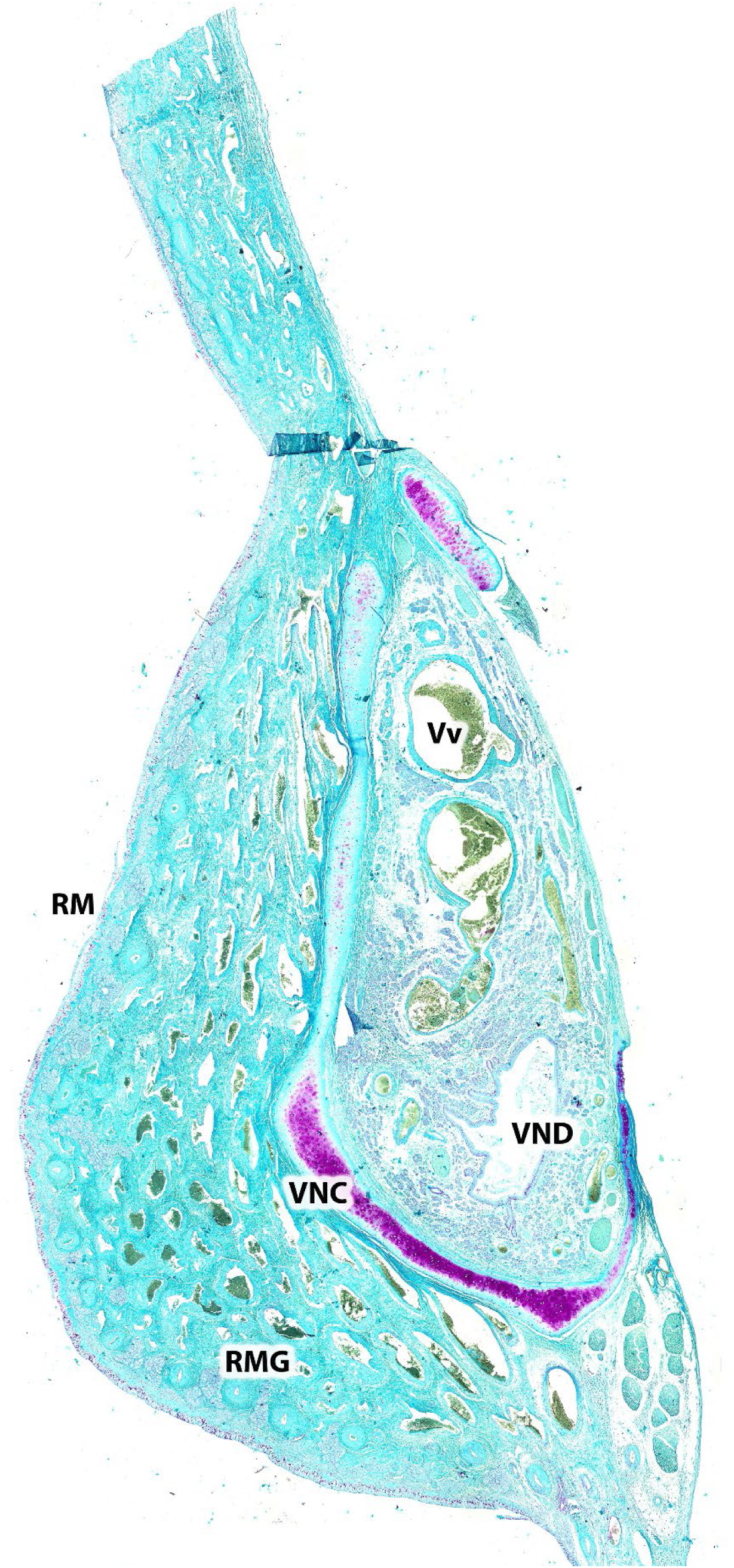
Histological image of the vomeronasal organ at a central level, stained with Gallego’s trichrome. The main components of the VNO are indicated: the vomeronasal cartilage (VNC), the vomeronasal duct (VND), the vascular pump (Vv)and the topographic relationship of the VNO with the respiratory mucosa of the nasal cavity (RM) which comprises glands (RMG) and blood vessels.

A serial transverse histological study of the VNO was conducted by analyzing representative sections across 14 serial rostrocaudal levels, aiming to characterize its morphology and longitudinal organization (Fig. 4). In the rostral region the initial configuration of the organ is observed, with the first signs of the developing vomeronasal duct. At these levels Gallego’s trichrome staining (Fig. 4A) and PAS (Fig. 4B) enabled the identification of extracellular matrix components and early secretory structures. Levels 2 to 14 are illustrated in Fig. 4C-Ñ. Across them, a progressive differentiation of the VNO architecture was noted. HE staining (Fig. 4C,D,F,H,J,K,M-O) highlighted cell nuclei and the organization of both sensory and respiratory epithelia, as well as the associated glands. Sections stained with Gallego’s trichrome (Fig. 4A,L) enhanced visualization of nerve fibers and connective tissue, while PAS staining (Fig.4B,G) emphasized the presence of carbohydrates in the glands and epithelial surfaces. AB-stained sections (Fig. 4E,I) revealed mucous components, particularly within the vomeronasal glands and surrounding mucosa.

**Figure 4.**
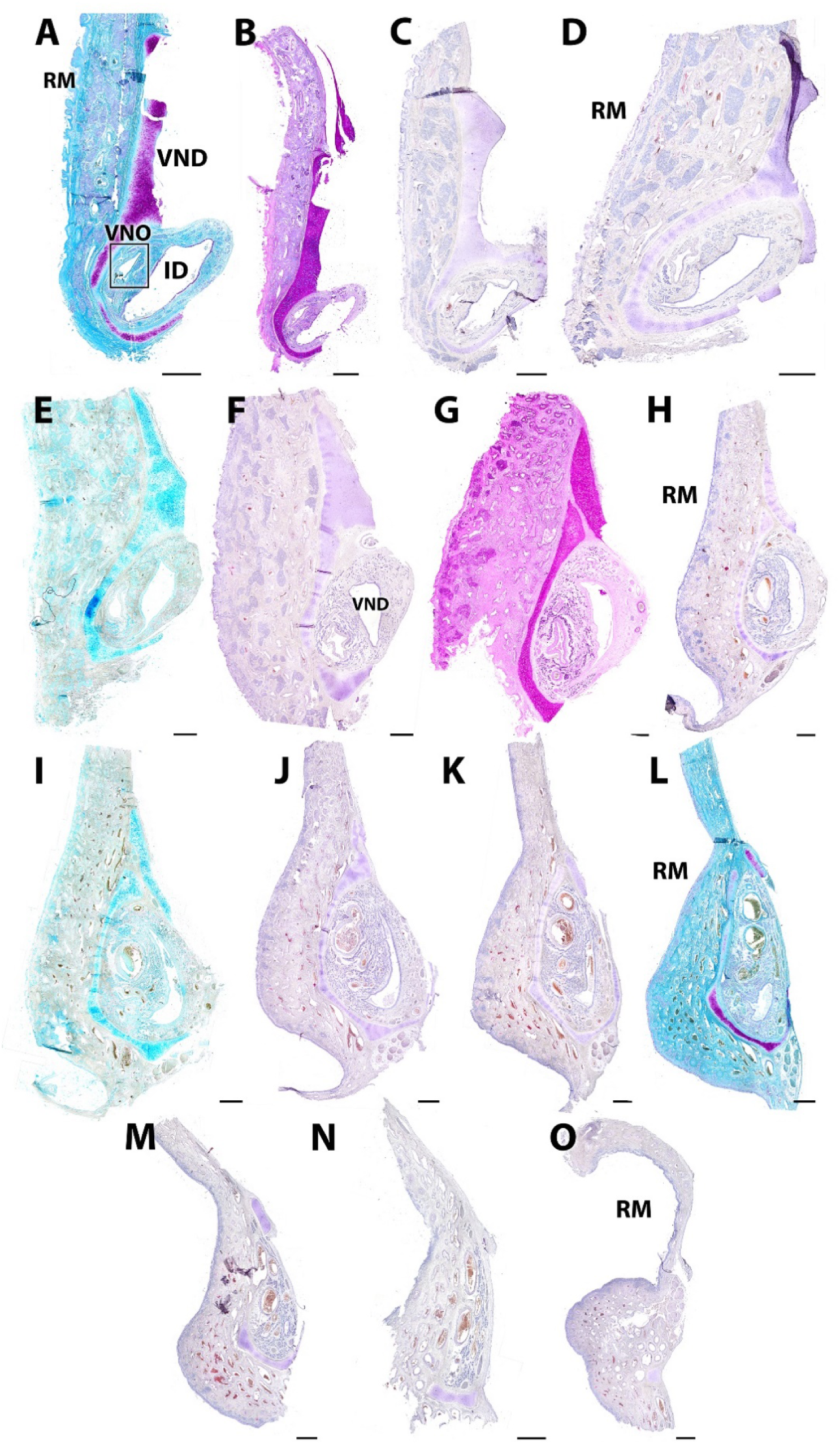
Transverse histological series of the wapiti VNO. This panel shows representative sections from the 14 rostrocaudal levels into which the wapiti’s VNO was divided for longitudinal analysis. Images A–B correspond to level 1, while the remaining 13 sections (C–Ñ) represent levels 2 through 14. Various histological stains were used throughout the study: Hematoxylin (C, D, F, H, J, K, M, N, O), Gallego’s trichrome (A–L), PAS (B, G), and Alcian blue (E,I). Scale bar: (A–Ñ) 1 mm.

Histological analysis of the vomeronasal duct revealed marked structural differences between the medial sensory (SE) and the lateral respiratory epithelia (RE), as assessed using various staining techniques. In sections from the caudal portion of the vomeronasal duct (Fig. 5A), the entire perimeter of the duct was lined with SE. Additionally, glands were observed completely surrounding the duct on both the medial and lateral sides. At a central level (Fig. 5B), the SE displayed a clearly layered organization: basal cells (1) were located at the base, followed by neuroepithelial cells (2), supporting cells (3), cellular processes (4), and microvilli projecting into the lumen (5). In contrast, the respiratory epithelium (Figures 5C-F) showed a pseudostratified organization over a well-defined basal cell layer. Within this epithelium, a central region with rounded nuclei at various heights was distinguished, along with an apical zone containing rounded nuclei and abundant cytoplasm. The underlying lamina propria exhibited a high density of glandular acini, some of which opened directly into the respiratory epithelium. Large-caliber venous vessels and atypical vascular arrangements were also evident.

**Figure 5.**
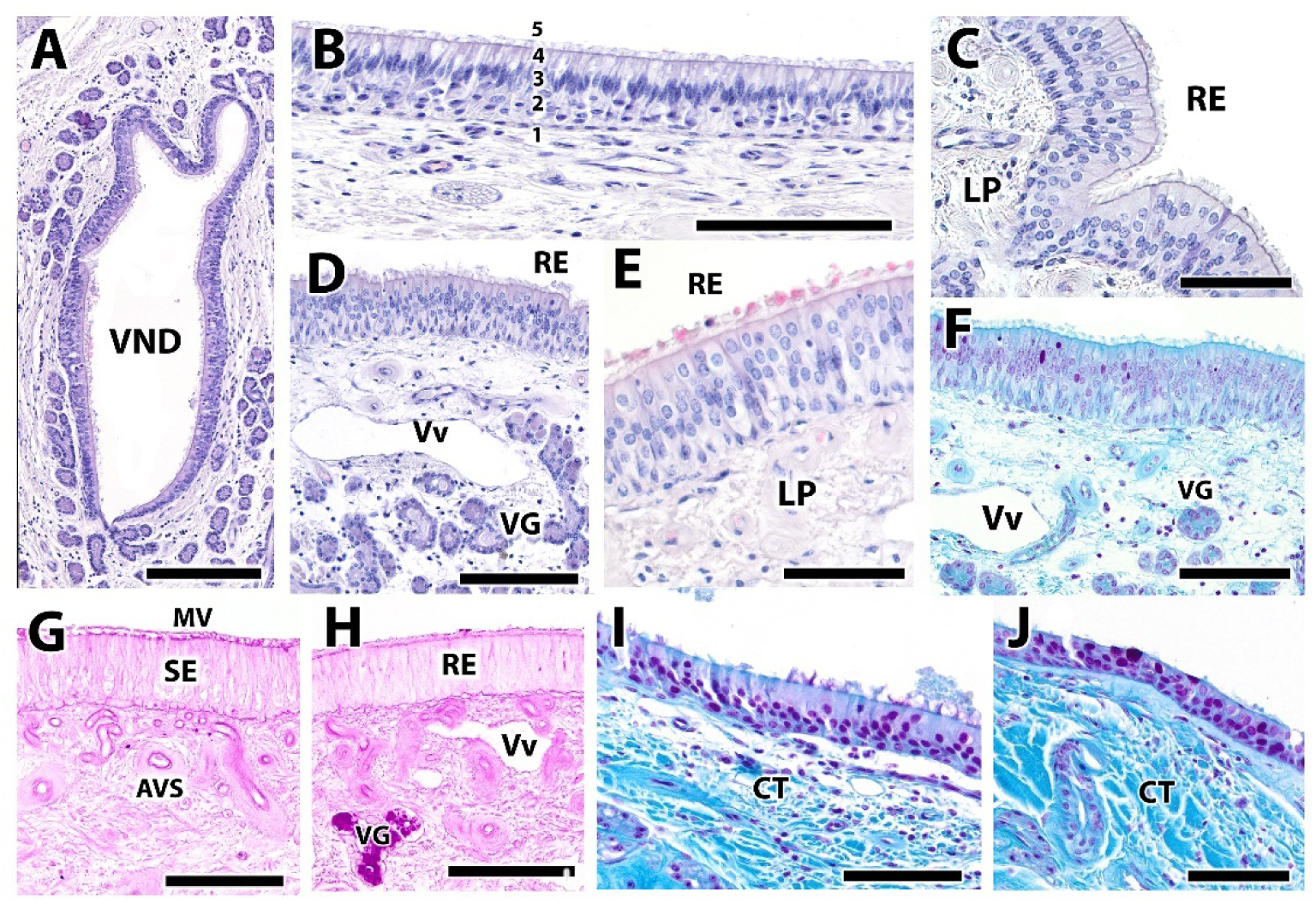
Histological comparison between the sensory and respiratory epithelia of the VNO in the wapiti, using various histological stains. A. General view of the caudal portion of the vomeronasal duct. At this level, the entire perimeter of the duct is lined by sensory epithelium. Glands are also seen surrounding the duct completely, both medially and laterally. B. Image of the sensory vomeronasal epithelium at a central level, showing its distinct layers numbered as follows: 1. Basal cells; 2. Neuroepithelial cells; 3. Supporting cells; 4. Cellular processes; 5. Microvilli. C–F. Histological images of the respiratory epithelium at various levels and stained with different techniques. All show a basal cell layer underlying a pseudostratified epithelium, which typically features a central region of rounded nuclei located at various heights and an apical layer of rounded nuclei with abundant cytoplasm. The lamina propria presents a high density of glandular acini, some of which open directly into the respiratory epithelium. Prominent venous vessels (Vv) and atypical vascular arrangements (AVA) are also present. G, H. PAS staining reveals a similar pattern in both sensory (G) and respiratory (H) epithelia. In both cases, the basal lamina stains intensely and glands are visible near the epithelium, especially in the respiratory type. No goblet cells were observed in either epithelium. I, J. Analysis of the most rostral levels of the VNO longitudinal series stained with Gallego’s trichrome. A markedly different epithelial architecture is observed: the epithelium, though stratified, is composed of tightly packed cells with small nuclei and projections extending into the lumen, particularly in the ventral zone (I), which appears more differentiated than the dorsal region (J). Scale bars: (A) 1 mm; (B, D, F–J) 200 μm; (C, E) 400 μm.

PAS staining (Fig. 5G,H) produced a similar staining pattern in both epithelial types, with intense labeling of the basal lamina and nearby glands, especially prominent in the respiratory epithelium. No goblet cells were identified in either type. Finally, the most rostral levels of the histological series, stained with Gallego’s trichrome (Fig. 5I,J), showed a distinct epithelial organization. Although stratified, the epithelium consisted of tightly packed cells with small nuclei and projections into the lumen, particularly in the ventral region (I), which appeared more differentiated than the dorsal area (J).

Histological analysis of the VNO (Fig. 6) allowed a clear identification of the main vascular and neural components involved in signal conduction within the parenchyma of the organ. At the most caudal level (Fig. 6A), a PAS-stained artery displayed an external wall composed of multiple concentric layers of elastin, consistent with the structure of large elastic vessels. Another artery, located at level 6 and stained with hematoxylin-eosin (Fig. 6B), was surrounded by abundant glandular tissue, indicating a close anatomical relationship between blood vessels and glands at this level of the VNO. A series of atypical vascular structures (Fig. 6C) were observed along the entire length of the organ, within the lamina propria. These profiles were characterized by thin, acellular PAS-negative walls and a central lumen lined by flat epithelium, distinguishing them morphologically from typical vessels.

**Figure 6.**
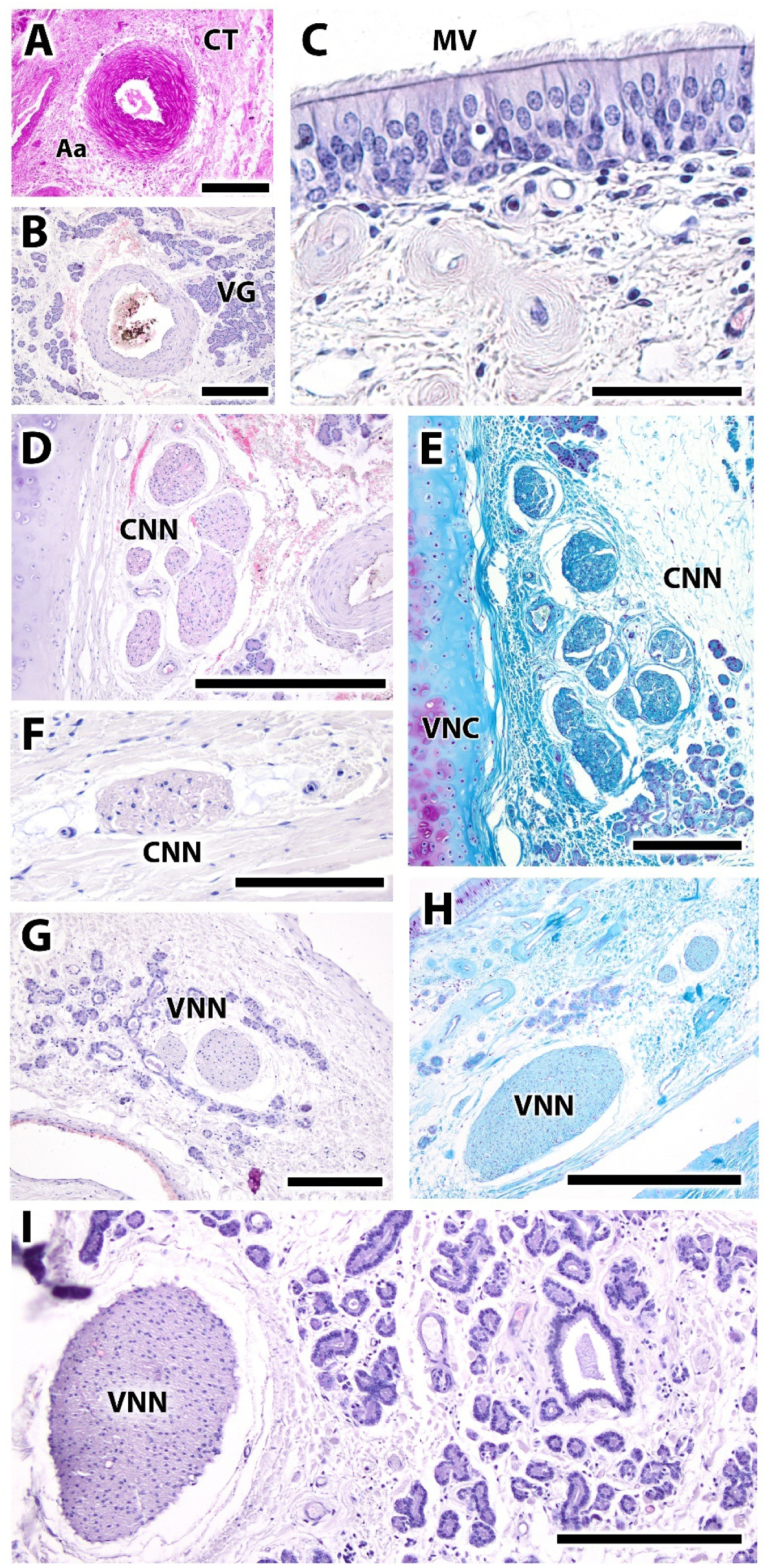
Composite image showing the conductive components (vascular and neural structures) of the wapiti VNO. A. Artery from the most caudal level of the VNO stained with PAS. The external wall is clearly composed of multiple concentric layers of elastin, characteristic of large-caliber elastic vessels. B. Artery from level 6 stained with hematoxylin-eosin (HE), surrounded by glandular tissue. C. Atypical vascular structures. Throughout the organ, the lamina propria contains a series of atypical vascular profiles characterized by an acellular PAS-negative wall and a central lumen lined with flat epithelium. D. Myelinated nerves. These are terminal branches of the caudal nasal nerve (CNN), located laterally within the parenchyma between blood vessels and glands. E. CNN stained with Gallego’s trichrome, showing the connective tissue surrounding the nerve fibers and their topographic relationship with the vomeronasal cartilage. F. Myelinated CNN stained with HE. These fibers correspond to vomeronasal axons originating from the sensory epithelium and located medially in the parenchyma, surrounded by glandular tissue. G. Vomeronasal nerves (VNN) are responsible for transmitting sensory information. H, I. VNNs as described above, stained with different techniques. Scale bars: (A, E, H) 500 μm; (C) 400 μm; (D, G) 250 μm; (F) 200 μm; (B, I) 125 μm.

Regarding the neural component, myelinated fibers belonging to the caudal nasal nerve (CNN) were observed in the lateral part of the parenchyma, situated between blood vessels and glands (Fig. 6D,F). Gallego’s trichrome staining allowed visualization of the connective tissue surrounding these fibers and their proximity to the vomeronasal cartilage (Fig. 6E). Vomeronasal axons (VNNs), originating from the sensory epithelium, were identified in the medial region of the parenchyma and also surrounded by glandular tissue (Fig. 6G).

These nerves are responsible for transmitting sensory information from the epithelium to the central nervous system. Figures 6H-I show similarly located VNNs stained with different histological techniques, highlighting their distribution in the caudal region of the organ and their close anatomical association with the surrounding glandular tissue.

Histological analysis of the glandular components associated with the VNO and the respiratory mucosa revealed a complex organization of glands with distinct biochemical composition and localization. Alcian blue (AB) staining, which detects acidic mucopolysaccharides, revealed intense positivity in the respiratory mucosa, where these substances form large alveolar-type glandular structures (Fig. 7A).

**Figure 7.**
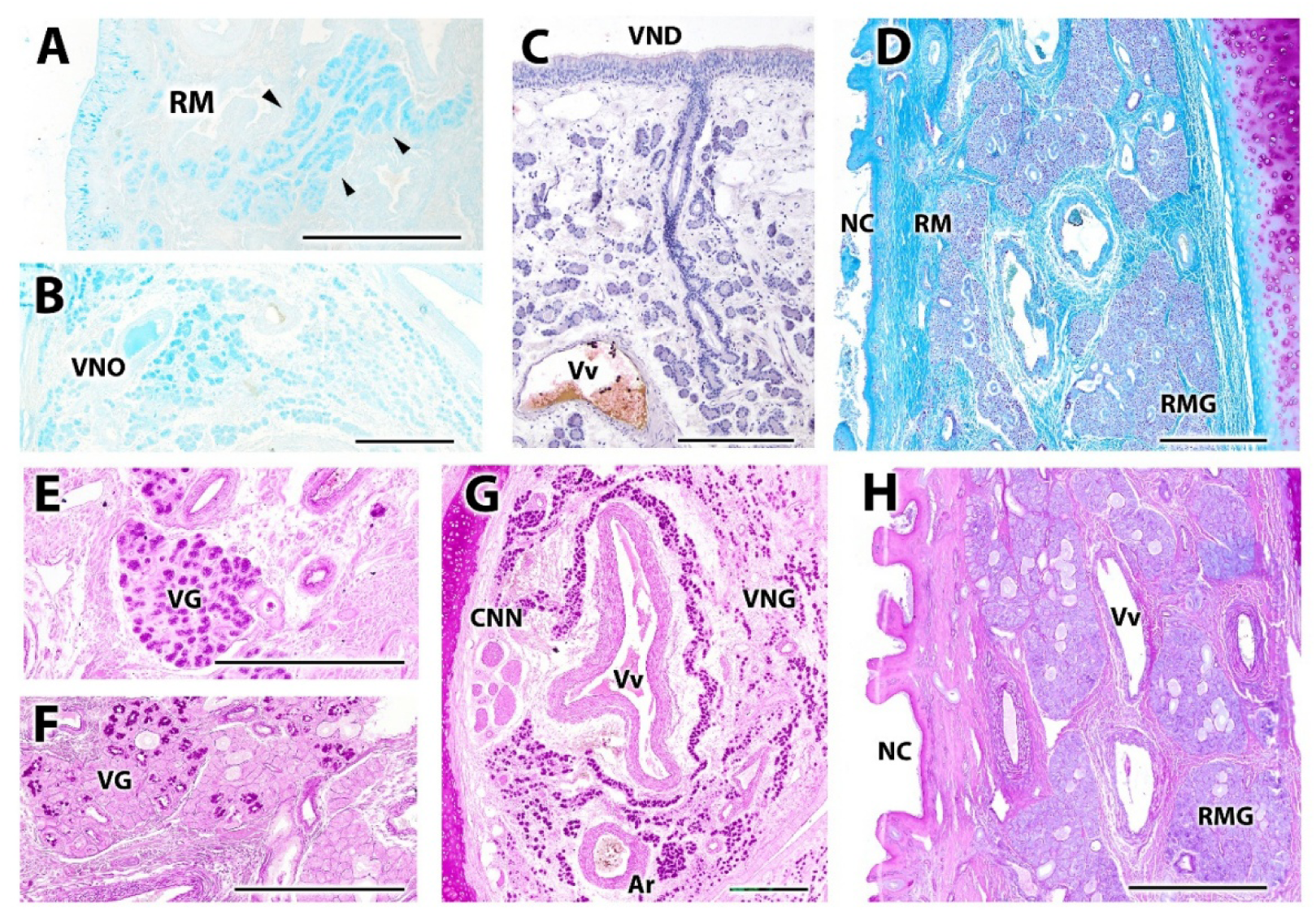
Histological images of the glandular component associated with the VNO and respiratory mucosa in the wapiti. A–B. Alcian blue (AB) staining highlights the presence of acidic mucopolysaccharides. These are found prominently in the respiratory mucosa forming large alveolar glands (A) but are also present in the VNO parenchyma (B). C. Hematoxylin-eosin (HE) staining reveals the development of the glandular component of the organ, showing alveolar structures connected to long ducts that open into the respiratory epithelium. D. Respiratory mucosa stained with Gallego’s trichrome, enabling visualization of both the glandular component and the surrounding connective tissue. E–G. PAS staining applied to the vomeronasal glands shows the presence of neutral mucopolysaccharides widely distributed among the glands. H. Respiratory mucosa stained with the combined PAS–AB technique reveals the presence of AB-positive glands in the respiratory mucosa, surrounded by PAS-positive connective tissue. Scale bar: (A–H) 500 μm.

Hematoxylin-eosin (HE) staining allowed for the observation of the structural development of the glandular system, revealing alveolar formations connected to long ducts that open directly into the respiratory epithelium (Fig. 7C). Gallego’s trichrome staining of the respiratory mucosa (Fig. 7D) facilitated the distinction between the glandular component and the surrounding connective tissue, showing a clear separation between these two tissue types. The PAS technique, which reveals neutral mucopolysaccharides, was applied to the vomeronasal glands (Fig. 7E-G), showing a broad distribution of these substances among the vomeronasal glands, suggesting an important neutral secretory activity. Finally, combined PAS–AB staining in the respiratory mucosa (Fig. 7H) confirmed the coexistence of AB-positive (acidic) glands surrounded by PAS-positive (neutral) connective tissue, demonstrating a complex interaction between both types of secretions within the glandular microenvironment of the respiratory mucosa.

Histological analysis of transverse sections of the wapiti’s palate mucosa revealed the anatomical arrangement of the incisive duct and its relationship with the VNO (Fig. 8). In sections taken at the mid-palatal level, a bilateral and symmetrical disposition of the incisive ducts and the rostral extremities of the vomeronasal organs was observed, both embedded within the thickness of the palatal mucosa. In this region, the ducts appeared well defined and clearly separated from each other, with no direct connection evident (Fig. 8A). It was also evident that the incisive papilla, vomeronasal cartilage, incisive bone, lamina propria, and palatal mucosa together form the structural foundation of the vomeronasal system in this area (Fig. 8A). This organization suggests a converging pathway toward the nasal cavity and a gradual functional transition along the rostrocaudal axis. The incisive papilla, in this region, appears as a central anatomical element mediating the connection between the oral cavity and the vomeronasal system. In more rostral sections, the complete fusion of the incisive duct with the incisive papilla was observed, along with the trajectory of the duct opening directly into the region of the VNO (Fig. 8B). At a more caudal level (Fig. 8C), the direct topographic relationship between the VNO and the incisive duct was confirmed, with both structures sharing a common parenchymal compartment.

**Figure 8.**
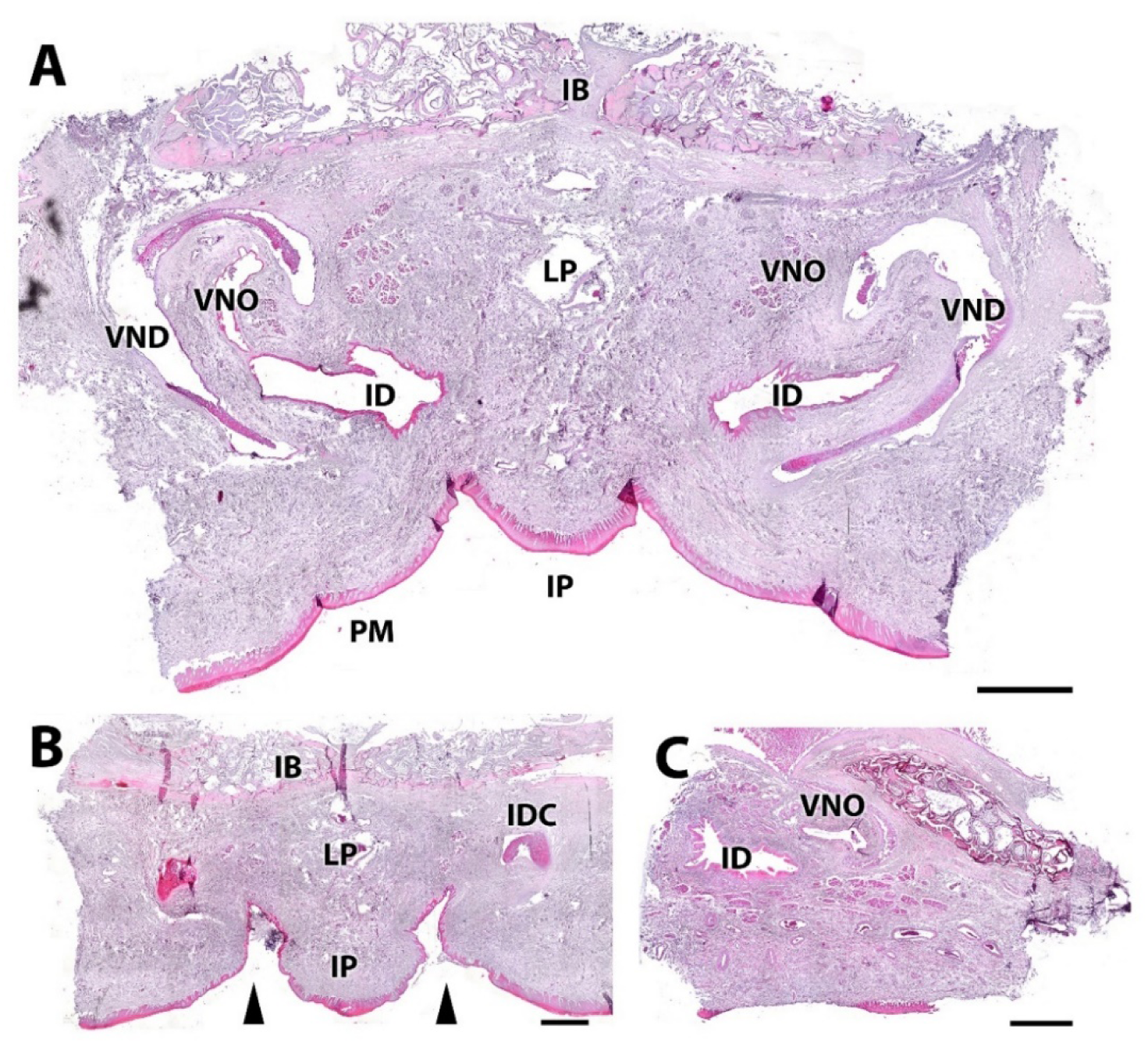
Histological organization of the palatal mucosa in the wapiti, showing the anatomical relationship between the incisive duct (ID) and the vomeronasal organ (VNO). A. Transverse section of the palate showing the bilateral and symmetrical arrangement of the VNO and the ID, both clearly separated on either side of the midline. Also visible are the incisive papilla (IP), vomeronasal cartilage (VNC), incisive bone (IB), lamina propria (LP), and palatal mucosa (PM). B. More rostral section where the ID and the VNO are positioned more closely together, beneath the IP (arrowheads), suggesting a converging pathway toward the nasal cavity. C. Caudal image showing the point at which the incisive duct runs parallel to the VNO, confirming a functional anatomical continuity between the oral cavity and the vomeronasal system. Scale bar:

#### Immunohistochemical study of the VNO

An immunohistochemical analysis of the vomeronasal duct epithelium in the wapiti (*Cervus canadensis*) was performed to characterize the distribution of various neuronal and glandular markers (Figs.9-10).

Immunolabeling with Gα0 revealed strong immunopositivity in the sensory epithelium, vomeronasal nerves, and associated glands, indicating expression in peripheral neural elements and secretory structures (Fig. 9A). Immunoreactivity for OMP was detected in both epithelial types (sensory and respiratory), as well as in nerves and glands, suggesting the presence of mature neurons across multiple regions of the organ (Fig. 9B). The PGP9.5 marker showed intense staining in the sensory epithelium, particularly in individual neurons, and exhibited strong labeling in vomeronasal nerves along with glandular positivity (Fig. 9C). Calretinin (CR) displayed immunoreactivity in the sensory epithelium and nerves, indicating association with differentiated neuronal populations (Fig. 9D). GAP43 expression was observed in nerves, glands, and the basal layer of the sensory epithelium— indicative of progenitor or differentiating cells—as well as in the ciliary layer of the respiratory epithelium, potentially reflecting neuronal plasticity in both regions (Fig. 9E). Finally, Gγ8 exhibited immunopositivity restricted to the sensory epithelium and vomeronasal nerves, indicating a preferential localization in vomeronasal system components (Fig. 9F).

**Figure 9.**
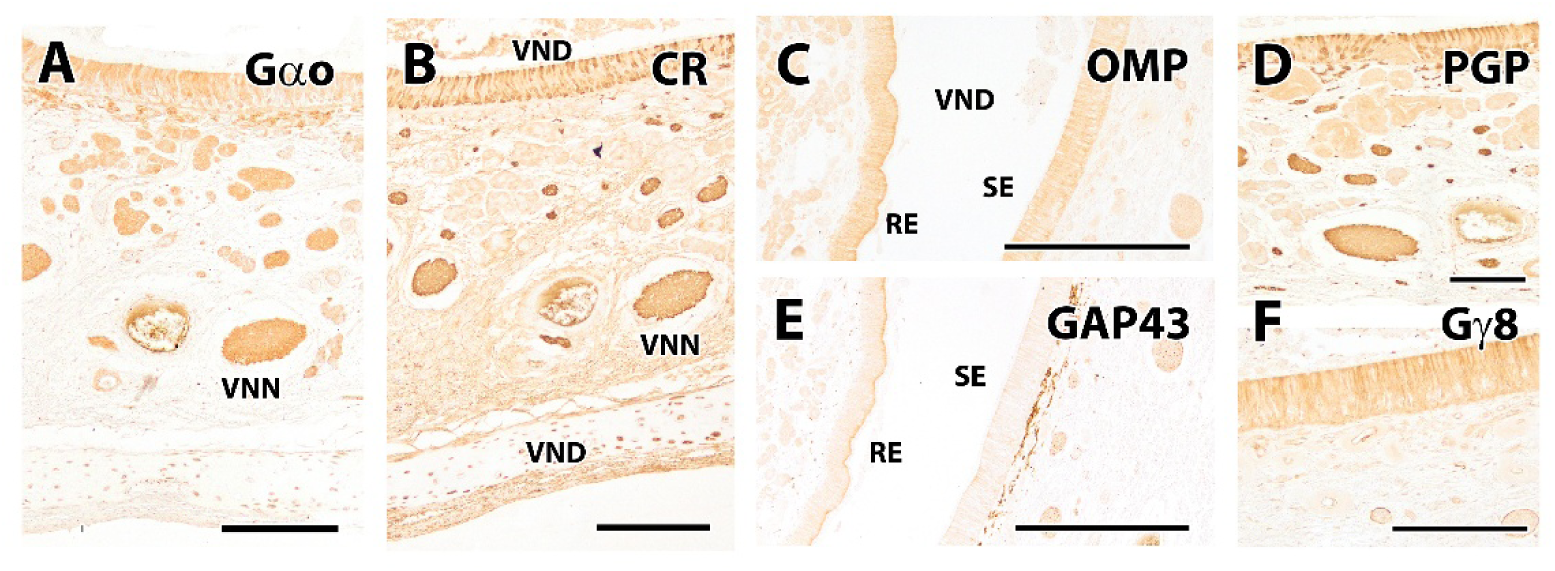
Immunohistochemical study of the vomeronasal duct epithelium (VDE) in the wapiti (*Cervus canadensis*). A. Gα0: strong immunoreactivity in the sensory epithelium (SE), vomeronasal nerves (VNN), and associated glands. B. Calretinin (CR): positive expression in the sensory epithelium, in neural pathways (VNN and VND), and in the glands. C. Olfactory Marker Protein (OMP): immunoreactivity observed in both the sensory and respiratory epithelium (RE), as well as in DVN nerve fibers. D. PGP9.5: robust labeling in the sensory epithelium, highlighting individual neurons; strong reactivity also noted in the vomeronasal nerves and associated glands. E. GAP43: expression in nerves, glands, the basal layer of the sensory epithelium, and the ciliary region of the respiratory epithelium. F. Gγ8: immunopositivity restricted to the sensory epithelium and underlying nerve fibers. Scale bars: (A, B, D, E) 500 μm; (C, F) 200 μm.

Anti-ROBO2 showed strong immunoreactivity in the sensory epithelium, with particularly intense expression in the basal layer, suggesting a role in signaling and epithelial organization during tissue maintenance or renewal (Fig. 10A). TUB3 displayed marked immunopositivity in both the sensory and respiratory epithelium, as well as in vomeronasal nerves, reflecting its structural role in neuronal components and differentiated epithelial cells (Fig. 10C). Substance P (SP) exhibited a homogeneous staining pattern across both epithelia and in the underlying glands, potentially indicating a shared role in sensory or secretory modulation in these regions (Fig. 10E).

**Figure 10.**
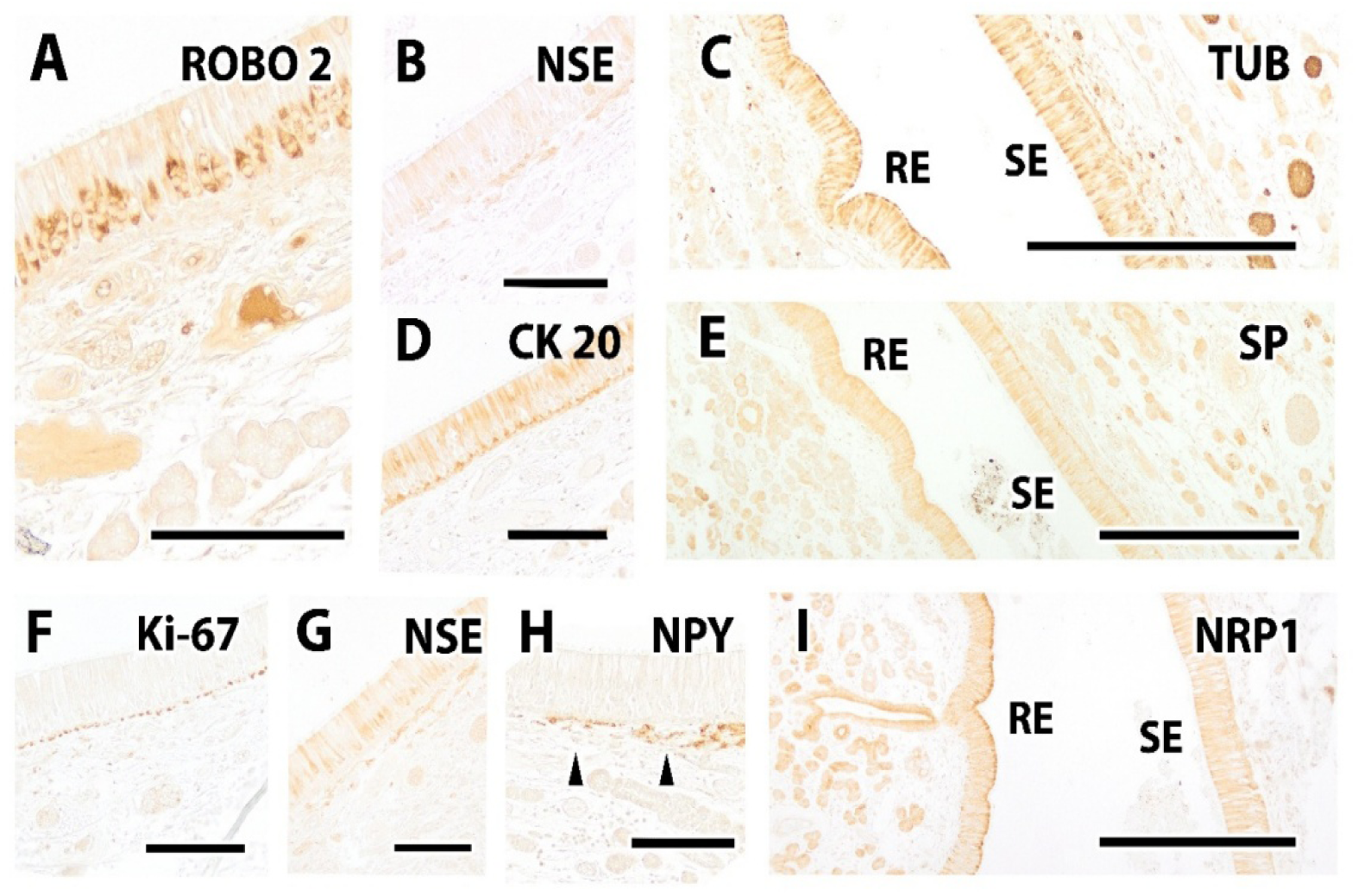
Immunohistochemical study of the vomeronasal duct epithelium in the wapiti. A. Anti-ROBO2 produced strong immunoreactivity in the sensory SE, with marked localization in the basal layer. B,G. Anti-NSE (neuron-specific enolase): weak labeling in the SE. C. TUB3 (β-tubulin III) showed a strong expression in both sensory and respiratory epithelia (RE), as well as in vomeronasal nerves. D. CK20 (cytokeratin 20): Its immunopositivity was restricted to the SE. E. SP (substance P): comparable reactivity in both epithelia and underlying glands. F. Ki-67: expression confined to the nuclei of basal epithelial cells, indicating active cell proliferation. H. NPY (neuropeptide Y) produced immunopositive labeling in the subepithelial layer immediately beneath the sensory epithelium (arrowheads). I. NRP1 (neuropilin-1): both epithelia showed immunoreactivity, with stronger signal intensity observed in the ciliary region of the RE. Scale bars: (A, D–H) 200 μm; (B, C, I) 200 μm.

CK20 was specifically expressed in the sensory epithelium, which may be associated with differentiated cellular subpopulations potentially involved in specialized functions (Fig. 10D). Ki-67, a proliferation marker, was confined to the nuclei of basal epithelial cells, indicating localized proliferative activity in the deeper epithelial layers (Fig. 10F). The enolase marker (EN) showed weak immunoreactivity in the sensory epithelium (Fig. 10B,G), while NPY was localized to the subepithelial layer immediately beneath the sensory epithelium, consistent with a local neuromodulatory or afferent-regulating function (Fig. 10H). Finally, NRP1 exhibited immunoreactivity in both the sensory and respiratory epithelium, with a notably higher intensity in the ciliary region of the respiratory epithelium, suggesting functional differences in the distribution of this molecule between the two epithelial types (Fig. 10I).

#### Lectin histochemical study of the VNO

A lectin histochemical study was conducted to characterize in detail the glycoconjugates expression in the tissue components of the VNO (Fig. 11). General views of the VNO revealed distinct staining patterns depending on the lectin used. DBA showed positivity on the surface of both epithelia, sensory and respiratory, especially in the ciliary region, as well as in glands of the VNO and respiratory mucosa (Fig. 11A). LEA produced a similar pattern, with intense labeling in the ciliary region, glands, and basal epithelial cells (Fig. 11B). SBA, applied to a more caudal region of the duct, also revealed positive labeling of the epithelial surface and glands (Fig. 11C). STL displayed a staining pattern closely resembling that of DBA (Fig. 11D).

**Figure 11.**
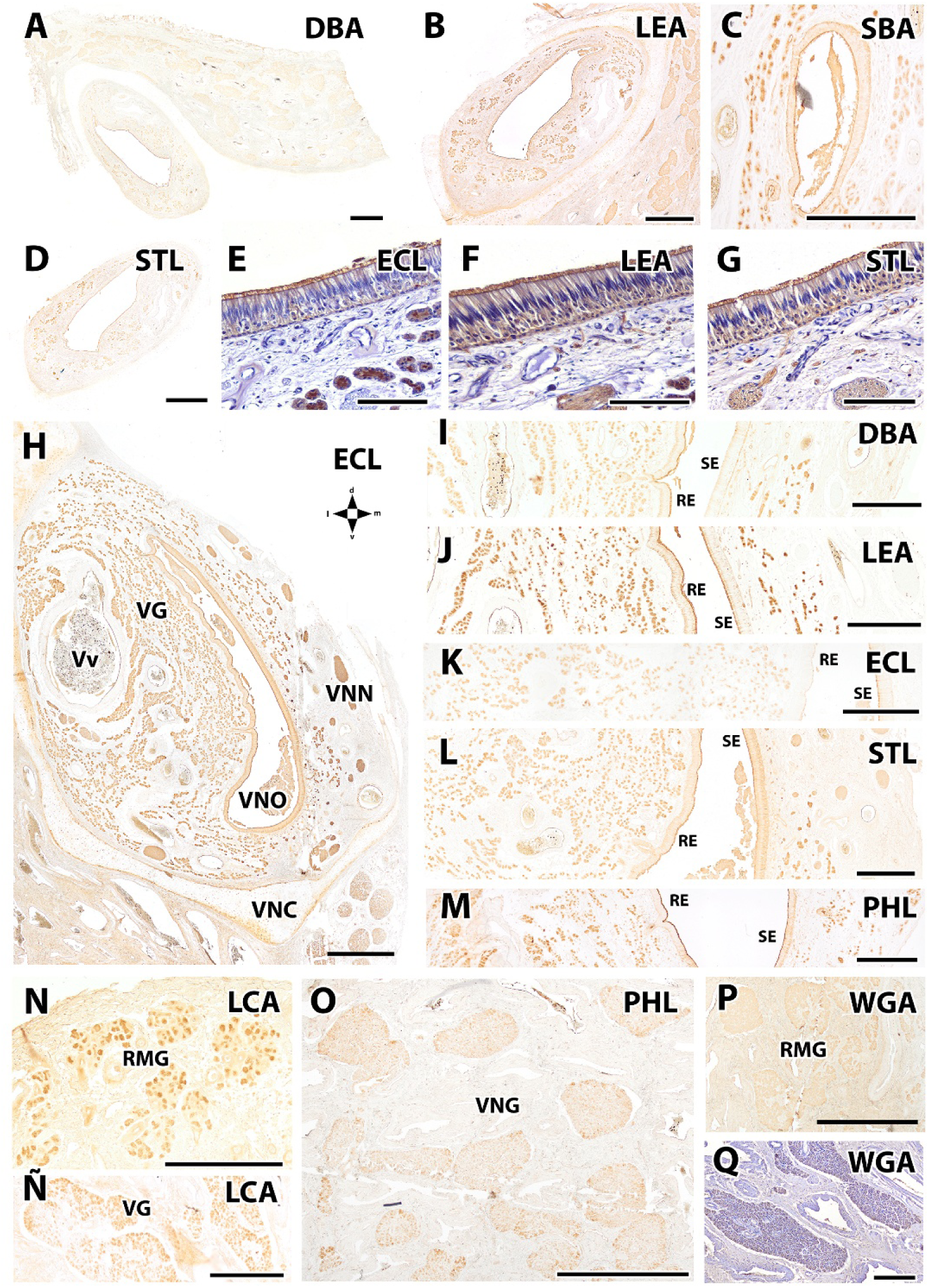
Lectin histochemistry of the wapiti VNO. A–D. General views of the VNO at different levels stained with DBA (A), LEA (B), SBA (C), and STL (D) lectins. Positivity is observed at the apical surface of both epithelia (especially in the ciliary region) and in glands from the RM and the VNO. E–G. Hematoxylin-eosin (HE) counterstained sections of the vomeronasal SE. E. ECL: strong reactivity in the ciliary region and glands, with weaker staining in the basal zone. F. LEA: intense labeling of cilia, basal cells, glands, and nerve bundles. G. STL: similar pattern to LEA. H. Overview of the VNO showing extensive reactivity in the ciliary region of the SE, vomeronasal nerves (VNN), vomeronasal glands (VG), and blood vessels (Vv). I–M. Comparative lectin binding in SE and RE. I. DBA: labeling in glands and ciliary surface of the RE, with no staining in the SE. J. LEA: similar staining pattern in both epithelia. K. ECL: positivity in the superficial SE; RE negative. Glandular reactivity is also present. L. STL: labeling in basal and apical regions of the SE, and only at the surface of the RE. Nerves and glands also positive. M. PHL: clear labeling in glands, SE, and respiratory epithelial cilia. N–Q. Lectin-binding profiles of VNG and respiratory mucosal glands (RMG). N. LCA: strong positivity in RMG with intense nuclear staining. Ñ. LCA: positive mucosal glands, with homogeneous morphology. O. PHL: positive labeling in mucosal glands. P. WGA: weak labeling in mucosal glands. Q. HE counterstain of the tissue shown in (P). Scale bars: (A, B, D, H, O, P) 1 mm; (C, I–M, N, Ñ, Q) 500 μm; (E–G) 200 μm.

HE-counterstained vomeronasal sensory epithelium sections revealed subcellular lectin localization. ECL showed strong expression in the ciliary region and glands, with weaker staining in the basal zone (Fig. 11E). A general view at a central level of the VNO revealed widespread reactivity in epithelia, nerves, and glands, with prominent positivity in the cilia (Fig. 11H). LEA revealed intense labeling in both basal cells and cilia, along with positivity in glands and nerve bundles (Fig. 11F). STL mirrored the pattern observed with LEA (Fig. 11G).

Comparative analysis between sensory and respiratory epithelia showed distinct lectin-binding profiles. DBA stained only the respiratory epithelium, with the labeling concentrated in the ciliary surface and glands (Fig. 11I). In contrast, LEA labeled both epithelia similarly, with reactivity in cilia and supporting cells (Fig. 11J). ECL presented a selective pattern: negative in the respiratory epithelium and positive in the superficial layer of the sensory epithelium, with additional expression in glands (Fig. 11K). Another lectin (STL) exhibited reactivity in both the basal and ciliary zones of the sensory epithelium and only at the surface of the respiratory epithelium. Nerves and glands were also positive (Fig. 11L). PHL labeled the glands, the sensory epithelium, and the ciliary surface of the respiratory epithelium (Fig. 11M).

The glandular component showed distinct lectin-binding affinities depending on origin. Vomeronasal glands displayed strong labeling with LCA, especially in the VNO parenchyma (Fig. 11N), while mucosal glands also exhibited LCA positivity but with more homogeneous morphology (Fig. 11Ñ). Mucosal glands were also labeled by PHL (Fig. 11O) and showed weaker reactivity with WGA (Fig. 11P). An enlarged HE-counterstained image of the latter allowed for clearer visualization of glandular architecture (Fig. 11Q).

#### Macroscopic study of the VNO

A macroscopic examination of the olfactory bulb in the wapiti (*Cervus canadensis*) was performed to characterize its general morphology, anatomical relationship with adjacent cranial structures, and to identify the potential position of the accessory olfactory bulb (Fig. 12). In the dorsal view of the skull (Fig. 12A), the dorsal surface of the cerebral hemispheres, the olfactory bulbs, and the ethmoidal fossa were clearly identified, allowing for spatial contextualization of the anterior olfactory system in relation to the brain. After removal and fixation of the olfactory bulb in Bouin’s solution (Fig. 12B-D), its shape and volume were more clearly defined, facilitating further histological processing. In the dorsal view of both the left and right OB (Fig. 12B,C), a distinct region marked with an asterisk was identified, presumably corresponding to the AOB based on its anatomical location. In the right OB it is possible to identify vomeronasal nerve fibers, further supporting the dorso-caudo-medial localization of the AOB in this species. Finally, in the anterior view of the cribriform plate of the ethmoid bone (Fig. 12E), the origin of the ethmoturbinals was evident, confirming the anatomical continuity between the nasal cavity and the base of the brain, as well as the expected trajectory of olfactory and vomeronasal nerves through the cribriform plate.

**Figure 12.**
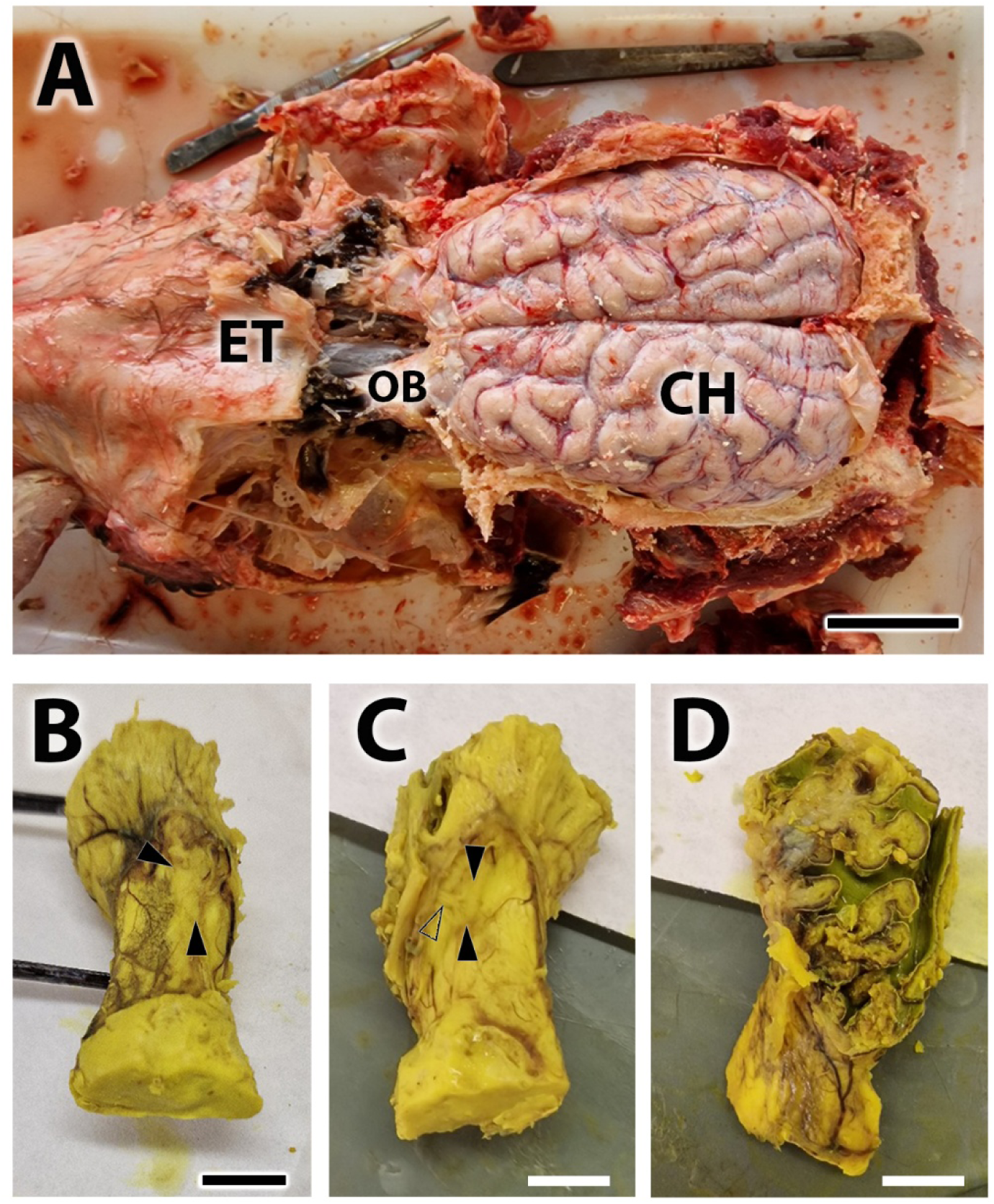
Macroscopic study of the olfactory bulb in the wapiti (Cervus canadensis). A. Dorsal view of the skull dissection, showing the dorsal surface of the cerebral hemispheres (CH), the olfactory bulb (OB), and the ethmoidal fossa (ET). B-D. OB after removal and fixation in Bouin’s solution. B. Dorsal view of the left OB, where the presumed location of the AOB can be observed (arrowheads). C. Dorsal view of the right OB, showing the presumptive AOB (arrowheads), with the arrival of the vomeronasal nerves (open arrowhead). D. Anterior view of the cribriform plate of the ethmoid bone, displaying the origin of the ethmoturbinals. Scale bars: (A) 5 cm; (B-D) 1 cm.

#### Microscopic study of the AOB

A serial histological examination of the anterior brain in the wapiti (*Cervus canadensis*) enabled characterization of the structure and topographic relationships of both the main olfactory bulb (MOB) and the accessory olfactory bulb (AOB).

In sagittal sections stained with hematoxylin-eosin (HE), the olfactory pathway from caudal to rostral was clearly delineated, including the olfactory peduncle (OP), the AOB, the arrival of vomeronasal nerves in close association with the ethmoidal artery, the olfactory limbus area, and the medial portion of the MOB (Fig. 13A). These structures define the full anterior trajectory of the olfactory tract in the brain.

**Figure 13.**
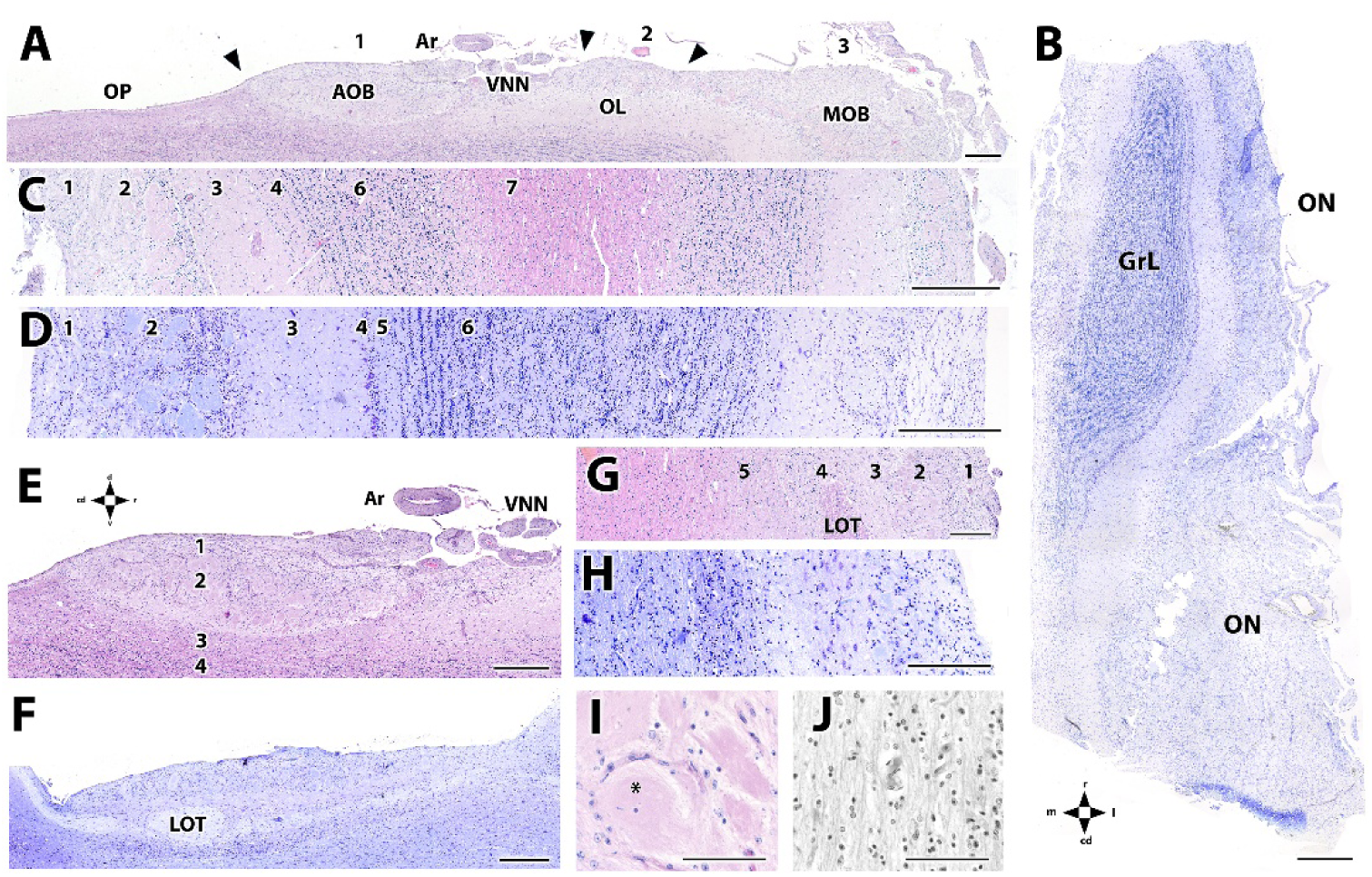
Histological study of the cellular organization of the AOB and MOB in the wapiti using hematoxylin-eosin and Nissl staining. A. Sagittal histological section of the medial margin of the olfactory bulb. From caudal to rostral, and delimited by arrowheads, the following structures are identified: the olfactory peduncle (OP), the AOB (1), the entry of the vomeronasal nerves (VNN) in direct association with the ethmoidal artery (Ar), the olfactory limbus (OL) area (2), and the MOB (3). Hematoxylin-eosin staining. B. Sagittal section of the MOB stained with Nissl showing clear lamination and prominent development of the olfactory nerve layer (ONL) and granule cell layer (GrL). C. Histological image of the MOB extending from the medial (left) to lateral (right) margin. A well-defined laminar organization is evident, consisting of: (1) olfactory nerve layer, (2) glomerular layer, (3) external plexiform layer, (4) mitral cell layer, (6) granule cell layer, and (7) white matter. D. Corresponding section stained with Nissl. E–F. Complete sagittal sections of the AOB stained with HE (E) and Nissl (F). G–H. Higher magnification views showing the laminar structure of the AOB: (1) vomeronasal nerve layer, (2) glomerular layer, (3) mitral-plexiform layer, (4) lateral olfactory tract (LOT), and (5) granule cell layer. I. Enlarged view of a glomerulus in the AOB. J. Lateral olfactory tract (LOT).

Nissl-stained sections of the MOB revealed a well-defined laminar architecture, with prominent development of the olfactory nerve layer (ONL) and granule cell layer (GrL), both notably expanded in this ungulate species (Fig. 13B). A full medial-to-lateral scan of the MOB allowed identification of all major histological layers: olfactory nerve layer, glomerular layer, external plexiform layer, mitral cell layer, granule cell layer, and white matter (Fig. 13C). This lamination was confirmed in consecutive panoramic sections stained with both HE and Nissl, which showed an almost identical distribution of the layers (Fig. 13C, D).

For the AOB, complete sagittal sections were processed with HE and Nissl staining (Figs. 13E,F), allowing direct comparison of structure and lamination with the MOB. The study revealed five distinct layers: (1) vomeronasal nerve layer, (2) glomerular layer, (3) mitral-plexiform layer, (4) lateral olfactory tract (LOT), and (5) granule cell layer (Figs. 13G,H). Mitral cells were more readily identified in Nissl-stained sections, whereas HE provided clearer distinction among layers. Detailed observation of individual glomeruli in both the MOB and AOB at higher magnification allowed the identification of periglomerular cells at the glomerular margins (Figs. 13H, I). Notably, the lateral olfactory tract (LOT) is particularly well developed in this species (Fig. 13J).

#### Immunohistochemical study of the AOB

A detailed immunohistochemical study of the accessory and main olfactory bulbs of the wapiti (*Cervus canadensis*) was performed to characterize the distribution of various proteins associated with neuronal signaling, development, and glial architecture (Figs. 14,15).

**Figure 14.**
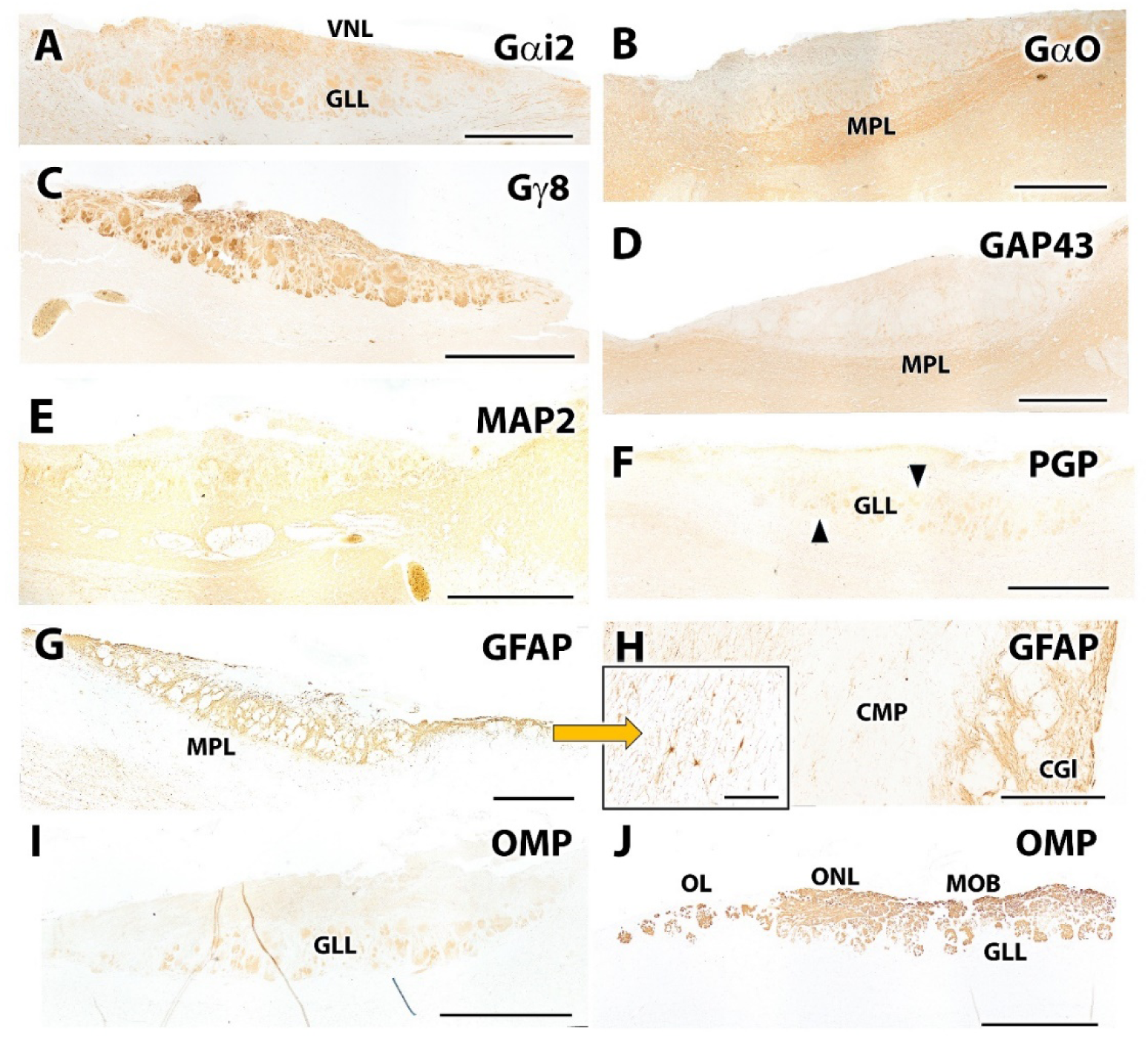
Immunohistochemical study of the accessory and main olfactory bulbs in the wapiti (I). A–C. Immunolabeling for G protein subunits. A. Anti-Gαi2 antibody shows moderate and widespread immunoreactivity in the superficial layers of the AOB, particularly in the vomeronasal nerve layer and the glomerular layer. B. Gα0 displays a complementary expression pattern, being mostly negative in superficial layers (except for occasional glomeruli) and strongly positive in deeper layers, mainly in the mitral-plexiform layer (MPL) and other regions of the telencephalon. C. Gγ8 shows superficial staining similar to Gαi2, but with greater intensity. D. GAP43 exhibits weak expression in the superficial layer but increased intensity in deeper layers, especially in the MPL; intermediate layers are negative. E. Anti-MAP2 shows intense labeling restricted to the deeper layers of the AOB and dendritic projections of mitral cells extending into the glomerular layer. The lateral olfactory tract (LOT) is immunonegative. F. PGP9.5 shows highly specific and intense immunoreactivity in the glomerular layer (arrowheads). G–H. GFAP immunostaining identifies the glial component, uniformly distributed among glomeruli of both the AOB and the main olfactory bulb (MOB). G. Positive GFAP labeling in the glomerular layer of the AOB; the MPL is immunonegative. H. Higher magnification of the GFAP-positive glomerular layer in the AOB, showing detailed views of glomeruli. The black inset highlights astrocytes (arrowhead) with visible processes. I–J. Immunohistochemistry for olfactory marker protein (OMP). I. OMP staining in the AOB shows positivity in the glomerular layer. J. Anti-OMP staining in the AOB and MOB shows positive labeling in the glomerular layer of both structures and in the olfactory limbus (OL). Unlike in the AOB, the superficial olfactory nerve layer (ONL) of the MOB shows strong immunopositivity. Scale bars: (A–J) 1 mm; (H inset) 200 μm.

**Figure 15.**
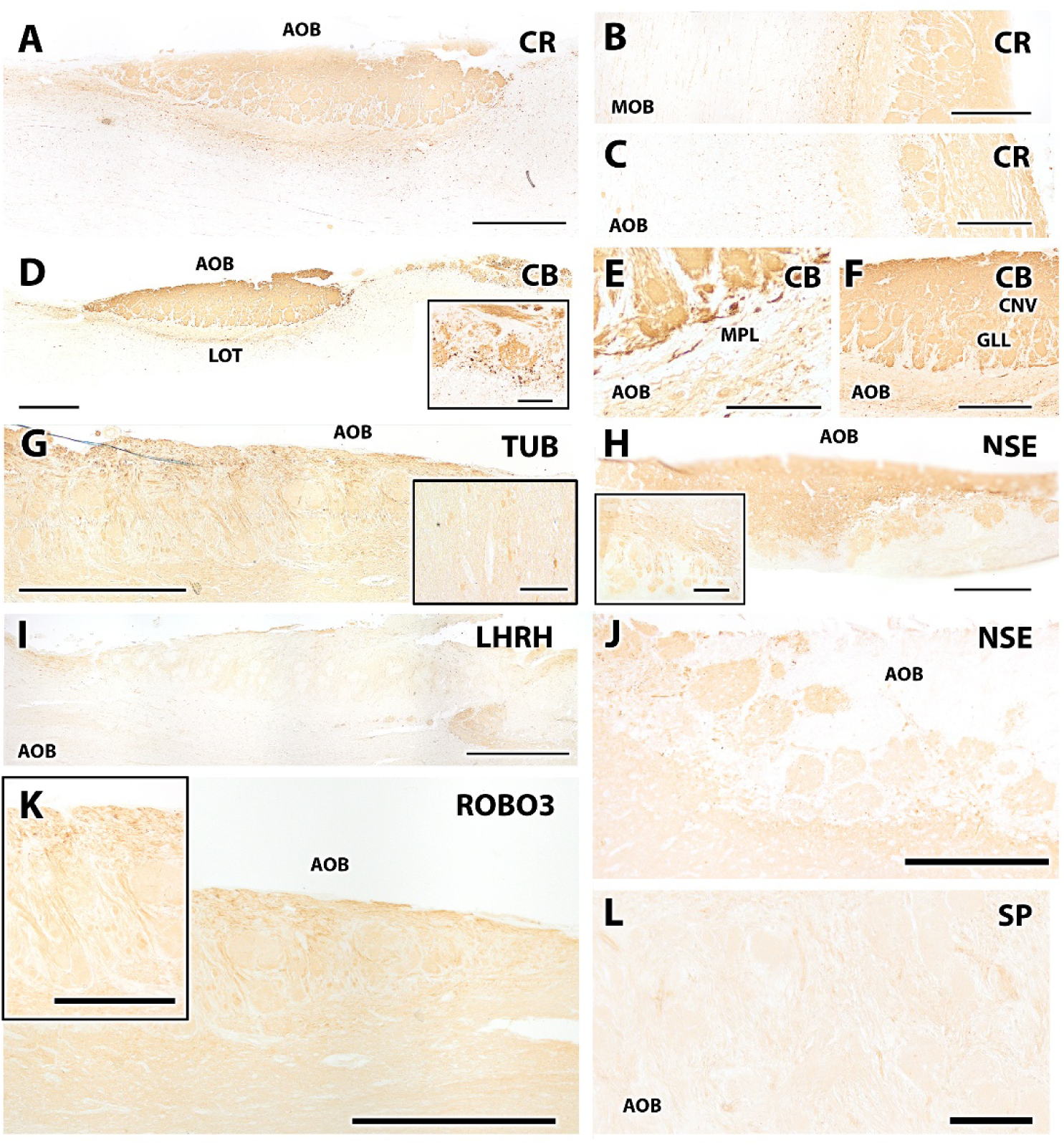
Immunohistochemical study of the accessory and main olfactory bulbs in the wapiti (II). A–C. Immunolabeling with anti-calretinin (CR). (A) Strong and widespread CR expression in the AOB, including the superficial nerve layer, glomerular layer, and mitral-plexiform layer. (B) CR immunopositivity in the MOB, observed in the superficial nerve layer, glomerular layer, and mitral layer. (C) Comparative view of the AOB, showing a labeling pattern equivalent to that of the MOB. **D–F.** Immunolabeling with anti-calbindin (CB). (D) Positive CB staining in most layers of the AOB and MOB, with absence of signal in the mitral-plexiform layer (AOB) and the mitral layer (MOB). The inset shows a magnified view of the MOB highlighting labeled glomeruli and periglomerular cells. (E–F) Higher magnifications of CB immunostaining in the AOB. Strong labeling in the glomerular layer is evident, although periglomerular cells remain unstained. The vomeronasal nerve layer (VNL) is intensely positive, while the plexiform layer (MPL) shows only faint reactivity. **G.** Tubulin expression in the AOB, showing homogeneous labeling across all layers. Inset shows a magnified area. **H.** Neuron-specific enolase (NSE) immunolabeling in the AOB, revealing superficial layer positivity, especially in glomeruli (inset). **I.** Immunoreactivity to GHRH in the AOB, highlighting immunopositive neuronal fibers. **J.** Additional NSE staining confirming strong signal in glomeruli and the mitral-plexiform layer of the AOB. **K.** Diffuse Robo3 expression in the AOB, with stronger signal in the neural component. **L.** Immunolabeling for substance P (SP) in the AOB, showing mild glial positivity. Scale bars: (A, D, G, H, I, K) 1 mm; (B, C, F, H inset, J, K inset) 500 μm.

Analysis of G protein subunits revealed complementary patterns between Gαi2 and Gα0. Gαi2 showed moderate and widespread immunoreactivity in the superficial layers of the AOB, particularly in the vomeronasal nerve layer (VNL) and the glomerular layer (GL), suggesting a role in early sensory signaling (Fig. 14A). In contrast, Gα0 was mainly expressed in deeper layers, showing minimal or absent staining in superficial zones except for isolated glomeruli, and strong positivity in the mitral-plexiform layer (MPL) and other telencephalic areas, indicating distinct functions at higher levels of sensory integration (Fig. 14B). Gy8 exhibited a superficial staining pattern similar to Gαi2 but with greater intensity, reinforcing its involvement in the sensory input zones of the AOB (Fig. 14C).

The plasticity-associated protein GAP43 showed weak staining in the most superficial layer but became progressively more intense in deeper regions, particularly in the MPL, with no signal detected in intermediate layers (Fig. 14D). The dendritic and somatic marker MAP2 displayed strong staining restricted to the deep layers of the AOB and dendritic projections toward the glomerular layer. The lateral olfactory tract (LOT) remained immunonegative, thus clearly defining its anatomical limits (Fig. 14E). PGP staining revealed highly specific immunoreactivity in the AOB glomerular layer, sharply delineating individual glomeruli (Fig. 14F). The glial profile, assessed by GFAP immunostaining, indicated positive labeling in the glomerular layer of both AOB and MOB, suggesting a uniform distribution of astrocytic elements (Fig. 14G). At higher magnification, astrocytes with their characteristic processes were clearly visible, and no labeling was observed in the mitral-plexiform layer (Fig. 14H). Finally, OMP staining showed clear positivity in the AOB glomerular layer (Fig. 14I). In the MOB and in the olfactory limbus (OL), intense immunopositivity was also observed in the superficial nerve layer (ONL), indicating the presence of mature olfactory neurons across all these regions (Fig. 14J).

Calretinin (CR) exhibited very intense and widespread immunoreactivity throughout the AOB, encompassing the superficial nerve layer, glomerular layer, and mitral-plexiform layer (Fig. 15A). This expression pattern was similarly observed in the MOB (Fig. 15B), and comparative views with the AOB confirmed the equivalent expression between both olfactory bulbs (Fig. 15C). Calbindin (CB) showed a more selective distribution. It was expressed in most layers of the AOB and MOB, with the exception of the mitral-plexiform layer in the AOB and the mitral layer in the MOB, which were devoid of labeling (Fig. 15D). In the MOB, intense labeling was observed in numerous glomeruli and periglomerular cells. In contrast, detailed images of the AOB (Fig. 15E, F) showed strong reactivity in the glomerular layer, while periglomerular cells were negative. The vomeronasal nerve layer (VNL) displayed marked immunoreactivity, while the mitral-plexiform layer exhibited weaker signal.

Tubulin immunostaining revealed a homogeneous distribution throughout the AOB without significant layer-specific differences (Fig. 15G). The neuron-specific enolase (NSE) marker showed predominant positivity in the superficial layers of the AOB, particularly within the glomeruli, as seen in the magnified view (Fig. 15H, J). LHRH immunostaining highlighted immunopositive neuronal fibers within the AOB (Fig. 15I), and supplementary NSE labeling further confirmed strong reactivity in both the glomerular and mitral-plexiform layers (Fig. 15J). Robo3 displayed a diffuse expression pattern within the AOB, with greater intensity in neural structures (Fig. 15K). Finally, substance P (SP) immunoreactivity was faint and restricted to the glial component of the AOB, suggesting a potential modulatory role at the neuroglial interface (Fig. 15L).

An immunohistochemical analysis using **free-floating sections** was performed to explore the glial and neuronal architecture of the AOB and the MOB. This method allowed the use of thicker sections, which enhanced three-dimensional tissue integrity and facilitated the visualization of complex neuronal structures and spatial relationships. A selected panel of markers was chosen to take advantage of these properties, including glial and neuronal proteins with known topographical specificity.

Immunolabeling with anti-GFAP revealed a well-defined glial distribution in both olfactory bulbs. In the MOB (Fig. 16-A), GFAP was strongly expressed between glomeruli and extended into the nerve layer, the granular layer, and the underlying white matter. In the AOB (Fig. 16-C), a similar labeling pattern was observed in the superficial layers, with clear delineation of individual glomeruli. The mitral-plexiform layer was notably GFAP-negative, confirming the regional specificity of astroglial localization.

**Figure 16.**
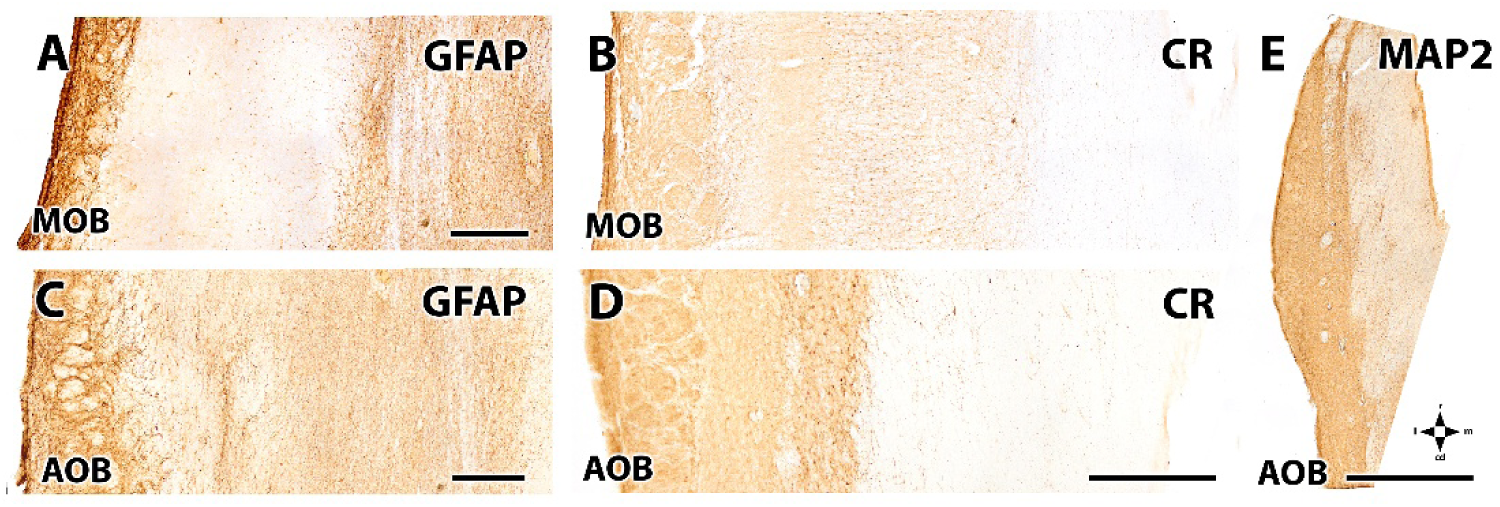
Free-floating immunohistochemical analysis of the accessory (AOB) and main olfactory bulbs (MOB) of the wapiti. A, C. GFAP immunolabeling. (A) GFAP staining in the MOB showing glial component between glomeruli, in the nerve layer, and in deeper layers including the granular layer and white matter. (C) GFAP staining in the AOB reveals a similar pattern with immunoreactivity in superficial layers, outlining the glomeruli; the mitral-plexiform layer is immunonegative. B, D. Calretinin (CR) immunolabeling. (B) Anti-CR staining in the MOB shows positive labeling across all layers. (D) CR staining in the AOB displays uniform positivity throughout the entire structure. E. MAP2 immunolabeling in the AOB shows strong and diffuse immunoreactivity across all layers. Scale bars: (A–D) 500 μm; (E) 1 mm.

Calretinin (CR) immunostaining demonstrated widespread neuronal labeling. In the MOB (Fig. 16-B), CR was present across all layers, consistent with its role in multiple neuronal subpopulations. Similarly, in the AOB (Fig. 16-D), strong immunoreactivity was detected throughout the structure, including superficial and deep layers, suggesting a uniform distribution of CR-expressing neurons. Finally, MAP2 immunolabeling in the AOB (Fig. 16-E) revealed intense and diffuse staining across the entire structure, indicative of its robust expression in neuronal somata and dendritic processes. The use of thick free-floating sections was particularly advantageous for visualizing MAP2-positive dendritic architecture in its full spatial extent.

#### Lectin histochemical study of the AOB

A lectin histochemical analysis was conducted to investigate the structural and glycoarchitectural organization of the accessory and main olfactory bulbs, using paraffin sections and a selected panel of plant lectins. This technique enabled the specific visualization of glycoconjugates in distinct olfactory layers and allowed comparison of glomerular architecture between both olfactory systems.

*Erythrina cristagalli* lectin (ECL) proved to be a specific marker of the glomerular layer in the AOB, with no labeling of the superficial nerve layer (Fig. 17A). ECL-positive glomeruli appeared as discrete, spherical structures with clearly defined boundaries, enabling accurate delineation of the sensory input region. In contrast, *Phaseolus vulgaris* lectin (PHL) displayed selective affinity for deeper layers of the AOB, with prominent labeling of the granular layer (Fig. 17B). Staining with *Solanum tuberosum* lectin (STL) revealed clear immunoreactivity in both the vomeronasal nerve layer and the glomerular layer of the AOB (Fig. 17C), highlighting the entry path of vomeronasal nerve fibers into the bulb. A similar pattern was observed with *Lycopersicon esculentum* agglutinin (LEA) (Fig. 17D).

**Figure 17.**
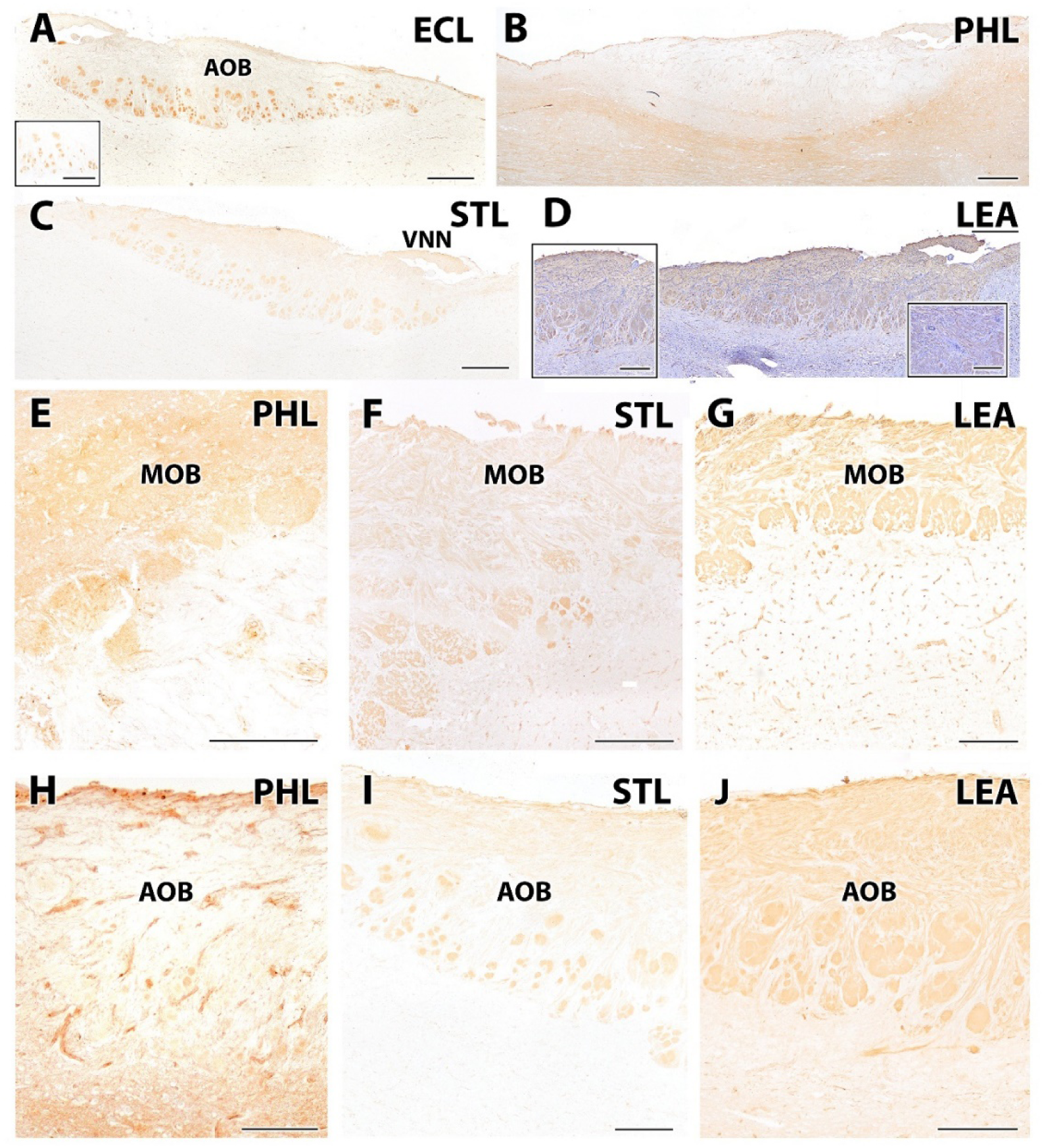
Lectin histochemical analysis of the accessory and main olfactory bulbs in the wapiti. A. *Erythrina cristagalli* lectin (ECL) shows specific labeling of the glomerular layer in the AOB, with no staining in the superficial nerve layer. Higher magnification (inset) reveals spherical, well-defined ECL-positive glomeruli. B. *Phaseolus vulgaris* lectin (PHL) preferentially stains the deep layers of the AOB, particularly the granular layer. C. *Solanum tuberosum* lectin (STL) labels both the vomeronasal nerve layer and the glomerular layer; the arrival of the vomeronasal nerve into the AOB is visible (arrowhead). D. *Lycopersicon esculentum* agglutinin (LEA) shows a similar pattern to STL, staining both the vomeronasal nerve layer and the glomerular layer. Inset shows higher magnification of these structures. E–J. Comparative analysis of MOB and AOB. E, H. PHL staining in the MOB (E) shows faint labeling, possibly restricted to capillary structures, while in the AOB (H) the staining is more intense in both the nerve and glomerular layers. F, G. STL (F) and LEA (G) staining in the MOB highlight intensely labeled glomeruli with spherical, homogeneous morphology and sharply defined boundaries. I, J. In the AOB, both STL (I) and LEA (J) label the glomerular layer; however, the glomeruli appear more irregular and less sharply delineated. Scale bars: (A–D) 500 μm; (Insets in A, D, and images E–J) 300 μm.

Comparative analysis between the MOB and AOB (Figs. 17E–J) demonstrated marked structural and staining differences. PHL labeling in the MOB (Fig. 17E) was faint and possibly restricted to capillary structures, whereas in the AOB (Fig. 17H) it showed intense labeling in the nerve and glomerular layers. Both STL and LEA produced robust glomerular labeling in the MOB (Figs. 17F–G), delineating well-defined, spherical glomeruli with homogeneous staining, indicative of a highly organized architecture.

Conversely, although the AOB also exhibited glomerular labeling with STL and LEA (Figs. 17I–J), the morphology of AOB glomeruli was more variable, with less defined boundaries and irregular shapes, suggesting underlying differences in sensory input organization and possibly functional integration between the AOB and MOB. These findings highlight the utility of lectin histochemistry for discriminating layer-specific glycoconjugate patterns and underscore the structural divergence between the two vomeronasal and main olfactory subsystems in this species.

#### Double immunohistochemistry for confocal fluorescence microscopy in free-floating sections

Confocal immunofluorescence revealed distinct organizational and neurochemical features of the main and accessory olfactory bulbs (MOB and AOB). Co-labeling with OMP and MAP2 (Fig. 18A,D) delineated the glomerular layer and neuronal architecture, showing a denser and more structured glomerular organization in the MOB compared to the AOB.

**Figure 18.**
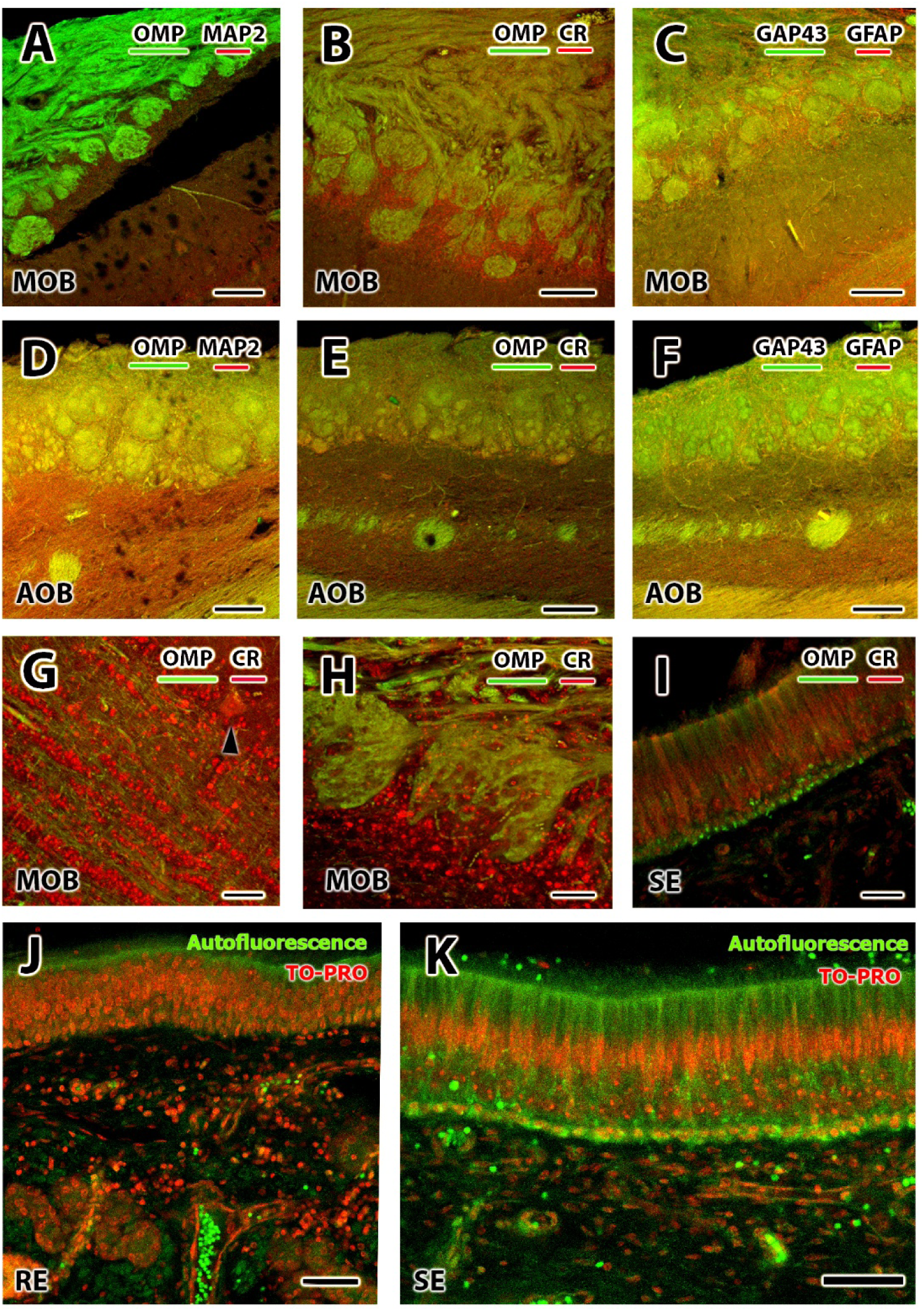
Immunofluorescence and autofluorescence labeling in the olfactory and vomeronasal systems of the wapiti. (A–F) Confocal double-labeling images of the main (MOB) and accessory olfactory bulbs (AOB). (A, D) Co-labeling with OMP (olfactory marker protein, green) and MAP2 (red) reveals the distinct glomerular organization of MOB (A) and AOB (D). (B, E, H) Double labeling with OMP (green) and calretinin (CR, red) in the MOB (B, H) and AOB (E) highlights periglomerular interneurons with strong CR expression, especially in MOB. Higher magnification in (H) emphasizes the periglomerular zone. (C, F) Co-labeling with GAP-43 (green) and GFAP (red) shows GAP-43-positive vomeronasal nerve fibers and GFAP-positive astrocytes within both MOB (C) and AOB (F). (G–I) Sensory epithelia: (G–H) Double labeling of OMP (green) and CR (red) in the MOB highlights granular cells, mitral cells (arrowheads), and periglomerular cells neurons. (I) CR+ neuroreceptor cells and OMP+ sustentacular and basal cells in the vomeronasal SE. (J–K) Autofluorescence (green) and nuclear counterstain TO-PRO (red) in respiratory (RE) and sensory (SE) epithelia. In SE (K), nuclear TO-PRO staining enables clear identification of neuroreceptor cell bodies, basal cells, and sustentacular cell somata and apical processes. Scale bars: (A–F) 250 μm; (G-K) 50 μm.

Double immunostaining with OMP and calretinin (Fig. 18B,E,H) highlighted periglomerular interneurons in both systems. These CR+ interneurons were particularly abundant and intensely labeled in the main olfactory bulb, where they formed a prominent ring around the glomeruli, as shown at higher magnification in panel H. In contrast, the AOB displayed fewer calretinin-positive cells.

Labeling with GAP-43 and GFAP (Fig. 18C,F) further differentiated nerve fiber and glial components. GAP-43-positive vomeronasal nerve fibers were concentrated in the superficial layers, while GFAP-positive astrocytes occupied deeper layers of both MOB and AOB.

In the sensory epithelia, co-localization of OMP and CR (Fig. 18G–I) marked receptor neurons and subpopulations of supporting and basal cells. In the sensory epithelium (SE), calretinin selectively labeled the apical dendritic layer and receptor neuron bodies, while OMP was distributed among sustentacular and basal cell populations.

Finally, TO-PRO nuclear staining combined with tissue autofluorescence (Fig. 18J– K) enabled morphological differentiation of epithelial cell types. In the respiratory epithelium (J), autofluorescence was confined to the apical layer, while TO-PRO highlighted the nuclei of all epithelial strata. In the sensory epithelium (K), the layered distribution of receptor neurons, basal cells, and sustentacular cells was clearly visible, with the latter showing distinct apical processes reaching the lumen.

## Discussion

The survival of organisms largely depends on effective chemical communication, particularly in ecological and reproductive contexts. Pheromones play a central role in this form of communication, as they are key regulators of sexual and social behaviors across all mammalian species (Burger, 2004). In order to fully understand how these substances influence behavior, a thorough and rigorous characterization of the systems responsible for their detection (the vomeronasal organ, VNO) and processing (the accessory olfactory bulb, AOB) is required. A crucial aspect that must be taken into account is the high degree of morphological and functional diversity not only among different vertebrate groups but also within them. This variability precludes interspecies extrapolation, highlighting the need for species-specific studies (Salazar and Sanchez Quinteiro, 2009).

This investigation focused on a wild North American ruminant species, the wapiti (*Cervus canadensis*), in which pheromonal cues are highly relevant due to the decisive role that sexual and social interactions play in its life history. The selection of this species was also motivated by the lack of comprehensive studies on wild ruminants. Therefore, we decided to focus our work on the wapiti. Given the absence of previous studies addressing these aspects in *Cervus canadensis*, we compared our findings with those reported in other species, primarily domestic ruminants.

Macroscopically, the VNO was easily identifiable on either side of the nasal septum due to its considerable size and length—features also observed in other ungulates such as the goat and domestic cattle (Park et al., 2013; Jang et al., 2022). Gross examination also allowed visualization of the vomeronasal nerves entering the AOB through its dorsomedial portion. This enabled us to confirm a direct connection between the VNNs and the AOB, unlike previous reports in some ruminants where such a connection was not demonstrated (Bertmar, 1981). Microscopically, we characterized the structural organization of the VNO and AOB. The VNO consists of two major components: cartilage and duct. The surrounding cartilage provides structural support and exhibits a J-shaped morphology similar to that seen in other mammals such as pigs and horses (Salazar et al., 1995). The inner part of the vomeronasal duct contains the neuroepithelium, essential for chemosensory reception.

Unlike most previous studies that typically examine a central segment of the organ—considered representative of the whole—or include only rostral and caudal views, our study applied a more exhaustive approach by examining the entire length of the organ. It was divided into 17 blocks, each 1 cm long, and both structural and neurochemical organization were analyzed for each segment. This allowed us to detect notable rostrocaudal anatomical and functional changes. For instance, sensory and respiratory epithelia were clearly differentiated only between blocks 3 and 10. Neither epithelium was fully developed at the rostral end, while the caudal regions displayed only sensory epithelium.

Between the vomeronasal duct and cartilage lies the parenchyma, a soft tissue region containing essential components for organ function. This area closely resembles that described in cattle (Salazar et al., 1997) and reindeer (Bertmar, 1981), with prominent venous sinuses, sparse arteries, unmyelinated vomeronasal nerves, and myelinated nasal caudal nerves. At the glandular level, our study revealed the presence of highly heterogeneous glands in the anterior region of the organ. The significance of this striking glandular pattern remains unknown.

Once the presence of a functional VNO was confirmed, we proceeded with a neurochemical analysis to characterize the molecules involved in chemosensory signal transduction. Immunohistochemical analysis included a broad panel of antibodies. Calcium-binding proteins such as calretinin (CR) and protein gene product 9.5 (PGP9.5) showed strong immunoreactivity. These markers are excellent indicators of vomeronasal activity and displayed expression patterns similar to those described in species like the fox (Ortiz-Leal et al., 2020) or wallaby (Torres et al., 2022), with positive labeling in both the sensory epithelium and vomeronasal nerves. The G protein subunits examined also showed positive expression. Their presence is associated with vomeronasal receptor activity, and the detection of Gα0 confirms the presence of V2R receptors in *Cervus canadensis*. These receptors are absent in many domestic species (Young and Trask, 2007), although studies in wild mammals (Ortiz-Leal et al., 2024) have also reported Gα0 expression. This may be linked to the importance of pheromonal communication in species with complex social hierarchies and parental care behaviors in natural environments. Furthermore, the expression of vomeronasal receptors varied along the length of the duct, suggesting that the organ may function as a chromatographic column. In this model, different molecules sampled by the VNO would travel through the duct according to their physicochemical properties and interactions with the mucus, thus allowing differential processing along the epithelial axis.

Other proteins, such as olfactory marker protein (OMP) and PGP9.5, exhibited expression patterns consistent with those found in the dama gazelle (Torres et al., 2023a) and the Korean roe deer (Park et al., 2014). This study also has included novel immunohistochemical markers in ungulates olfactory system, such as ROBO2 (Prince et al., 2009) and CK20 (Khan et al., 2021). Particularly noteworthy was the result obtained with Ki-67, which stained exclusively basal cells, indicating a potential role in early neuronal differentiation (Naik et al., 2020).

Lectin histochemistry was conducted to characterize the glycoconjugate composition of VNO. This information is useful for identifying distinct epithelial domains, assessing functional compartmentalization, and uncovering species-specific patterns of glycosylation that may underlie differential pheromone detection and signal transduction mechanisms (Keller et al., 2022). LEA lectin showed positivity in both the sensory and olfactory epithelia, indicating its involvement in both the main and accessory olfactory pathways. This pattern is consistent with findings in other ruminants such as sheep (Salazar et al., 2000; Ibrahim, 2018), as is also the case with PHL lectin (Yang et al., 2021). These results support the notion that conserved glycosylation profiles contribute to the functional integration of chemosensory inputs across olfactory subsystems in ruminants.

Following the VNO analysis, we examined the AOB, the main center for vomeronasal signal integration. As in the VNO, the organization of the AOB varies significantly across species. This variability ranges from highly laminated structures in rodents (Suárez et al., 2011) and marsupials (Schneider, 2011) to its complete absence or extreme reduction in species such as the dog (Miodonski, 1968).

In *Cervus canadensis*, the AOB was large enough to be observed macroscopically. Microscopically, it displayed clear lamination with all the layers typically described, although the mitral–plexiform layer was less developed than that seen in mice, rats and rabbits (Larriva-Sahd, 2008; Villamayor et al., 2020). A novel finding in this species was the identification of extensive white matter tracts traversing the structure, consistent with the lateral olfactory tract described in some mammals, although they appeared as discrete, individualized bundles with an atypical caliber when compared to those observed in other mammalian groups (Switzer, III. et al., 1980).

Immunohistochemical analysis of the AOB revealed positive expression of the G protein subunits Gαi2 and Gγ8 in the superficial layers of the AOB, including the vomeronasal and glomerular nerve fibers. This pattern indicates active expression of the V1R receptor family (Jia and Halpern, 1996). In contrast, Gα0 showed a complementary distribution, being localized to the mitral–plexiform layer (Takigami et al., 2000). The absence of Gα0 labeling in the AOB suggests that Gα0-positive neurons identified in the vomeronasal epithelium may not project to the AOB. Their projection target remains unknown; however, based on studies conducted in the fox (Ortiz-Leal et al., 2023), it can be hypothesized that these neurons project to the transitional zone between the main and accessory olfactory bulbs, where specific chemosensory signals might be processed. Histological analysis of serial AOB sections confirmed that this transition area is characterized by smaller, less defined glomeruli with principal cells arranged peripherally. OMP expression was detected in both the main and accessory olfactory bulbs, consistent with its presence in both VNO epithelia. Similarly, CR and CB were expressed in both pathways, matching descriptions in other placental mammals such as the capybara (Torres et al., 2020). To our knowledge, no previous studies have reported free-floating immunohistochemistry in the AOB of ruminants. Establishing this technique allowed us to clearly distinguish between glial (GFAP-positive) and glomerular (CB-positive) components and to validate the paraffin-based immunolabeling results. GAP43 was negative in both the AOB and the VNO, suggesting that this protein may not be involved in the accessory pathway.

Lectin staining in the AOB revealed that PHL was specifically expressed in the mitral–plexiform layer and absent in other regions of the AOB, suggesting a role in synaptic or transport processes, or a selective expression by specific neuronal subtypes in that layer. As observed in the VNO, LEA was positive in both the main and accessory bulbs, supporting the consistency of the overall vomeronasal system profile in this species.

The results of this study confirm that the vomeronasal system in Cervus canadensis is highly developed, both anatomically and neurochemically. The rostrocaudal differentiation of the VNO, the complex glandular architecture, and the organized expression of immunohistochemical and lectin markers all point to a functionally active and specialized system. This structural and molecular complexity suggests that pheromones play a critical role in regulating social and reproductive behavior in this wild species. Consequently, the wapiti represents a valuable model for exploring chemical communication mechanisms in ungulates and for understanding the evolutionary and ecological implications of these sensory systems in natural environments.

## AUTHOR CONTRIBUTIONS

Conceptualization, A.M.A., E.V.C., P.S.Q., and I.O.L.; Methodology, A.M.A., G.G.H., E.V.C., P.S.Q., and I.O.L.; Investigation, A.M.A., G.G.H., E.V.C., J.L.R., O.S.M., P.S.Q. and I.O.L.; Resources, E.V.C., J.L.R., and P.S.Q.; Writing – Original Draft Preparation, A.M.A.; Writing – Review & Editing, A.M.A., P.S.Q, and I.O.L.; Supervision, P.S.Q. and I.O.L.; Project Administration, P.S.Q. and I.O.L.; Funding Acquisition, P.S.Q.

## INSTITUTIONAL REVIEW BOARD STATEMENT

Not applicable, as all the animals employed in this study died by natural causes.

## INFORMED CONSENT STATEMENT

Not applicable, as this research did not involve any humans.

## DATA AVAILABILITY STATEMENT

All relevant data are within the manuscript and are fully available without restriction.

